# Competition for endothelial cell polarity drives vascular morphogenesis

**DOI:** 10.1101/2021.11.23.469704

**Authors:** Pedro Barbacena, Maria Dominguez-Cejudo, Catarina G. Fonseca, Manuel Gómez-González, Laura M. Faure, Georgia Zarkada, Andreia A. Pena, Anna Pezzarossa, Daniela Ramalho, Ylenia Giarratano, Marie Ouarné, David Barata, Isabela Fortunato, Lenka H. Misiková, Ian Mauldin, Yulia Carvalho, Xavier Trepat, Pere Roca-Cusachs, Anne Eichmann, Miguel O. Bernabeu, Cláudio A. Franco

## Abstract

Blood vessel formation generates unique vascular patterns in each individual. The principles governing the apparent stochasticity of this process remain to be elucidated. Using mathematical methods, we find that the transition between two fundamental vascular morphogenetic programs – sprouting angiogenesis and vascular remodeling – is established by a shift on collective front-rear polarity of endothelial cells. We demonstrate that the competition between biochemical (VEGFA) and mechanical (blood flow-induced shear stress) cues controls this collective polarity shift. Shear stress increases tension at focal adhesions overriding VEGFA-driven collective polarization, which relies on tension at adherens junctions. We propose that vascular morphogenetic cues compete to regulate individual cell polarity and migration through tension shifts that translates into tissue-level emergent behaviors, ultimately leading to uniquely organized vascular patterns.

## Introduction

The blood vascular network is a branched system irrigating all organs in vertebrates, which is fundamental for embryogenesis, physiology and healing. Dysfunction of this network is associated with multiple diseases, including cancer progression, diabetic retinopathies, or arteriovenous malformations (Potente and Makinen, 2017). Major axial vessels are stereotypical and are formed through vasculogenesis (Potente and Makinen, 2017). Yet, expansion of this early embryonic network through angiogenesis (Potente and Makinen, 2017), seems stochastic because it generates vasculatures with unique patterns, which can be used for biometric identification (Hartung et al., 2012). Angiogenesis involves two distinct morphogenetic processes: *i*) sprouting angiogenesis, which relies on chemokines, such as vascular endothelial growth factor A (VEGFA). This process expands pre-existing networks through proliferation, migration and anastomosis of endothelial cells (ECs), and forms immature networks. *ii*) vascular remodeling, which relies on blood flow-induced shear stress. This process converts immature networks, generated by sprouting angiogenesis, into hierarchical vascular networks, requiring vessel pruning, arteriovenous differentiation and vessel specialization (Korn and Augustin, 2015; Potente and Makinen, 2017). How ECs shift between these two morphogenetic processes, and which principles govern the formation of well-organized, yet unique, networks remain outstanding questions in vascular biology.

Sprouting and remodeling morphogenetic programs involve dynamic coordination of cell polarity and migration (Fonseca et al., 2020; Korn and Augustin, 2015), which are regulated by cell-level and tissue-level integration of chemical and mechanical signals. For instance, sprouting angiogenesis involves adherens junction (AJ)-mediated mechanotransduction regulating collective migration downstream of VEGFA stimuli (Cao et al., 2017; Carvalho et al., 2019; Friedl and Mayor, 2017; Hayer et al., 2016). In addition, ECs sense and respond to blood flow-induced shear stress by aligning, polarizing and migrating against the flow direction, a phenomenon known as flow-migration coupling (Franco et al., 2015; Kwon et al., 2016; Park et al., 2021; Tanaka et al., 2021; Tzima et al., 2001). Despite the well-known crosstalk between VEGFA and shear stress to regulate EC sprouting capacity (Chouinard-Pelletier et al., 2013; Ghaffari et al., 2015; Song and Munn, 2011), there is a lack of understanding of how these inputs are integrated at the cell level to shape the vascular network at the tissue scale *in vivo*. Here, we investigated how chemical (VEGFA) and mechanical (shear stress) cues interact to promote changes in cellular behaviors underlying the transition between sprouting-to-remodeling morphogenetic programs, which we termed as the S>R transition.

## Results

### Endothelial polarity patterns define the S>R transition zone

To understand how chemical (VEGFA) and mechanical (shear stress) cues establish the S>R transition, we analyzed the mouse retina vasculature, where VEGFA and shear stress are in opposite gradients (Figure 1A). To identify the S>R transition, we first measured 7 vessel morphometric features (Figure S1A) in 100*μ*m-wide bins along the retina (Figure 1A), following the sprouting front-to-optic nerve axis (Figure 1B). Principal component analysis (PCA) classified each bin based on phenotypic similarity (see Methods). Each bin point was assigned to one of the two expected biological classes (sprouting or remodeling), using the k-means clustering algorithm (Figure 1C). The frequency distribution of bins from class 1 (sprouting) and class 2 (remodeling) along the sprouting front-to-optic nerve axis determined a transition at 300-400*μ*m from the sprouting front (Figure 1D), corresponding to a shift in the predominance of class 1 to class 2 bins. Thus, this unbiased quantitative method based on morphometric features captured a transition in vascular organization, the morphological S>R transition zone.

**Figure 1.**
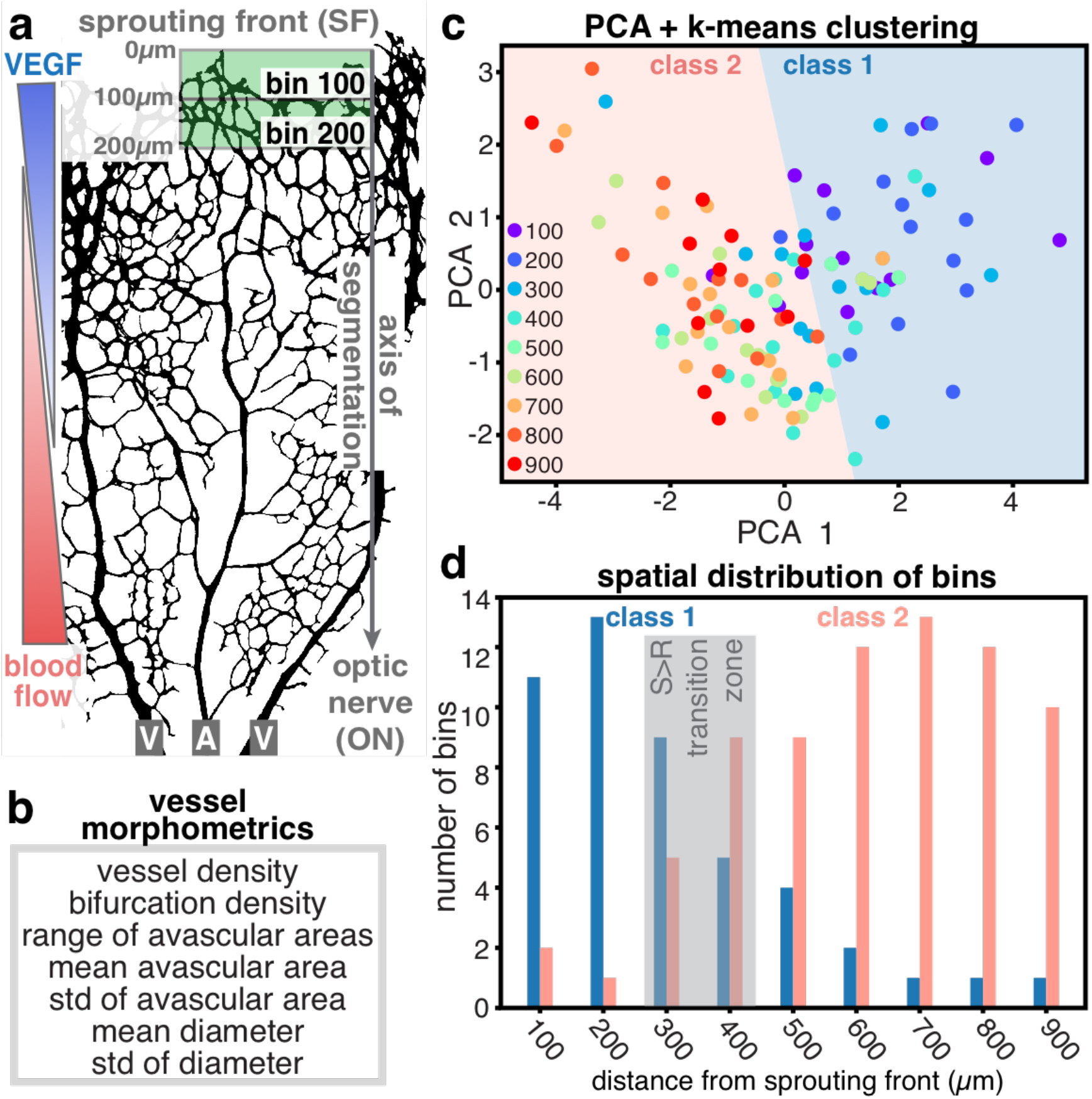
Vessel morphometrics define the morphological S>R transition. **A**, Schematic of the sprouting front-to-optic nerve (SF>ON) segmentation axis of the retinal vasculature, and its relationship to VEGFA and shear stress gradients. Segmentation bins 100 and 200 (width 100*μ*m) are depicted as green rectangles. V, vein; A, artery. **B**, Set of parameters quantified in each bin, termed as vessel morphometrics. **C**, Principal component analysis (PCA) of vessel morphometrics in wild-type vascular networks followed by k-means clustering. Each dot represents one bin in one retina, color-coded for the corresponding bin number. K-means clustering identifies 2 classes of objects, class 1 (sprouting) and class 2 (remodeling). N=14 retinas. **D**, Distribution of the number of class 1 or class 2 bins along the SF>ON axis. Grey rectangle defines the morphological S>R transition zone.

Next, we investigated how this morphological transition could be explained at the cellular level, based on chemical and mechanical cues. Sprouting angiogenesis and vascular remodeling rely on EC migration (Fonseca et al., 2020; Korn and Augustin, 2015; Potente and Makinen, 2017). EC polarity is a readout for cell migration, and both VEGFA and blood flow induce EC polarization and migration in zebrafish and mouse (Carvalho et al., 2019; Franco et al., 2015; Kwon et al., 2016). Therefore, we hypothesized that changes in EC polarity could underlie the morphological S>R transition. To investigate this interaction, we analyzed the angle between the nucleus-to-Golgi polarity axis and the sprouting front edge (K-angle), or the simulated blood flow direction (F-angle), for each EC (Figure 2A, Figure S1B). The analysis was performed using PolNet (Bernabeu et al., 2018), on the same bins (100*μ*m wide) used for vessel morphometrics (Figure 1A). At the sprouting front (bin 100*μ*m), K-angle centered at ~90° (towards the VEGFA gradient) with a narrow scattering (±14.8°), whilst F-angle had a higher dispersion (mean 197.3° ±31.7), indicating a poor influence of flow (Figure 2B). At 400 *μ*m away from the sprouting front, blood flow gained a strong influence on EC polarity, with an F-angle centered at 180° with a very narrow dispersion compared to bin 100 *μ*m (~178.8° ±4.2; p= 0.0012 for Levene test for unequal variances). This was associated with a significant shift for the mean K-angle from ~90° (100*μ*m) to ~270° (400*μ*m) (p= 0.0016 for Kruskal-Wallis ANOVA test), representing an inversion of polarity from “towards” to “against” the sprouting front (Figure 2B and Figure S1C). From K- and F-angles, we calculated K- and F-indexes, standing for chemokine-dependent or flow-dependent polarity indexes, respectively (Figure 2C, Figure S1D). K- and F-indexes quantify the robustness of EC polarity towards (positive values) or against (negative values) a given polarity cue (details in Methods). Notably, K- and F-indexes are anti-correlated (R^2^= −0.838), highlighting an interdependency between the two polarity cues (Figure S1E). Closer to the sprouting front, the absolute value of K-index was higher than F-index, demonstrating the dominance of the chemokine stimulus (0.22 ±0.03 vs −0.09 ±0.07 at 100*μ*m, respectively) (Figure 2C). Yet, away from the sprouting front region, K- and F-indexes inverted, with K-index becoming negative and lower than F-index (−0.21 ±0.04 vs 0.32 ±0.05 at 400*μ*m, respectively), demonstrating the dominance of flow in determining EC polarity (Figure 2D). Thus, based on the mathematical description of EC polarity, we defined the cellular S>R transition as a 100*μ*m width (bin width) region centered at the point where the mean F-index intersect the mean of K-index (Figure 2D). The global analysis locates the cellular S>R transition at 230 ±50*μ*m from the sprouting front (Figure 2D), whilst analysis per retina sets it at 203.3 ±23.0*μ*m (Figure 2E). Anatomical mapping places the cellular S>R transition just ahead of the artery tip in the mouse retina (Figure 2F). Overall, we propose a novel system-level mathematical description of EC polarity to predict the S>R transition; that the cellular S>R transition zone (180-280*μ*m) (Figure 2D) precedes the morphological S>R transition zone (300-400*μ*m) (Figure 1D); and that the polarity landscape of ECs is dominated by the effect of shear stress, revealing a narrow region of influence for chemokine signaling.

**Figure 2.**
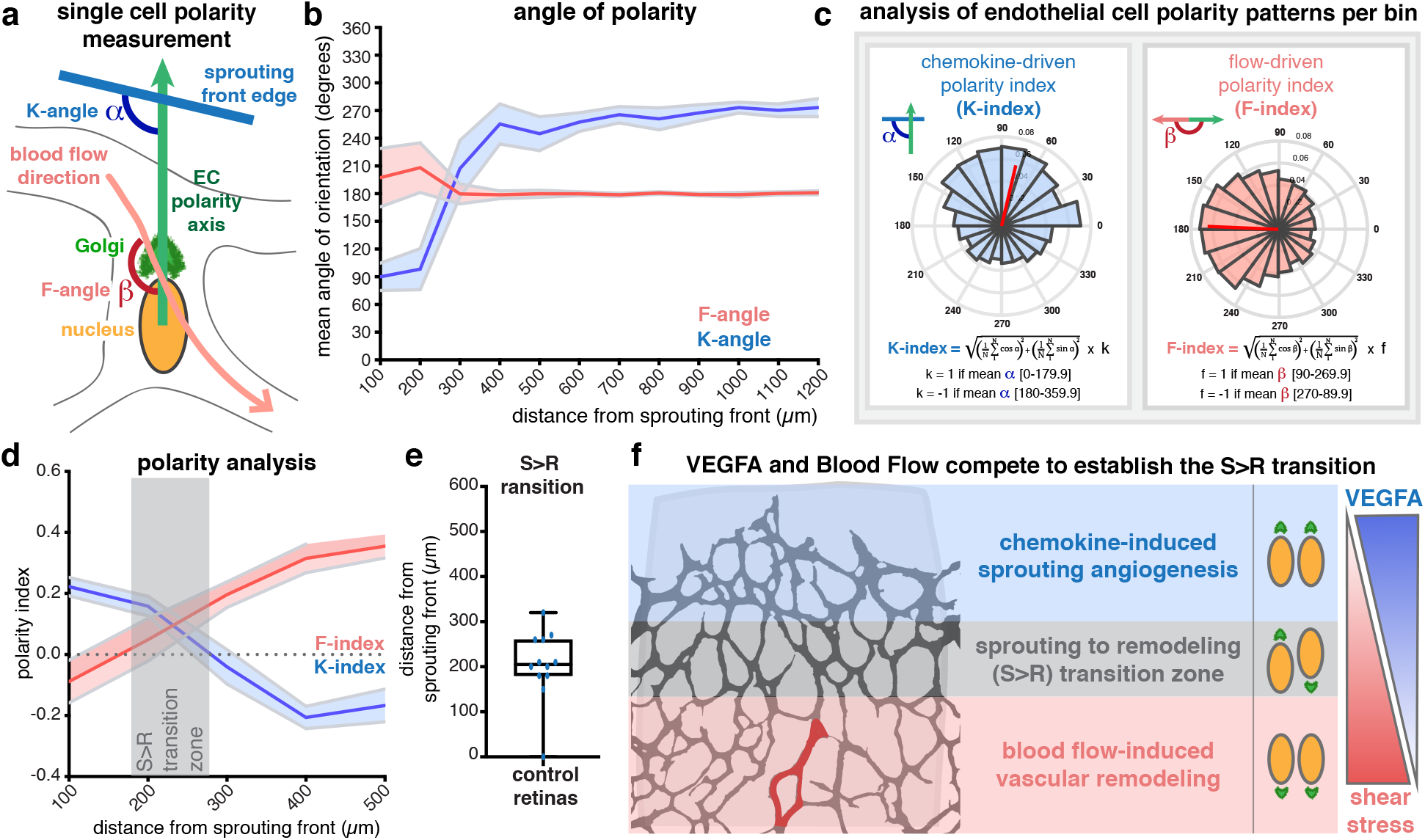
Collective EC polarity patterns establish the cellular S>R transition. **A**, Representation of the calculation method for chemokine-induced (K) and flow-induced (F) angles based on individual EC polarity (nucleus-to-Golgi vector) axis. **B**, Distribution of K-angles (blue) and F-angles (red) along the SF>ON axis. Solid line represents mean, and light area represents SEM. **C**, Calculation of K- and F-indexes, based on K- or F-angles, respectively. **D**, Distribution of K-(blue) and F-indexes (red) within the 500*μ*m from the SF (dashed box in Figure S1C). Solid line represents mean, and light area represents SEM. Dashed black line represents random polarity. Grey rectangle defines the cellular S>R transition zone. **E**, Box plot for the cellular S>R transition in each retina. **F**, Morphological annotation of the cellular S>R transition in a mouse retina. Red vessel corresponds to the tip of the retinal artery. N=11 retinas.

### Competition between VEGFA and shear stress levels define the S>R transition zone

Our analysis of the S>R transition predicts that increasing levels of VEGFA or reducing shear stress should promote sprouting angiogenesis. Conversely, reduction of VEGFA levels or increasing shear stress levels should promote vascular remodeling. Thus, we manipulated blood flow or VEGFA to disturb the S>R transition. First, we treated animals with captopril (an angiotensin-converting enzyme inhibitor) to reduce blood flow, or angiotensin-II to increase blood flow (Nehme et al., 2019). Captopril treatment led to a significant increase in vessel density and in the number of endothelial tip cells, associated with sprouting angiogenesis (Figure 3A, Figure S2A-G), which correlated with a spatial shift on the cellular S>R transition, from ~230*μ*m to ~325*μ*m (Figure 3B). Angiotensin-II treatment decreased vascular density and the number of tip cells, abolishing the S>R transition zone (Figure 3C,D, Figure S2A-G). Next, we manipulated VEGFA levels in the mouse retina by intraocular injection of VEGFA or sFLT1 (a VEGFA inhibitor) (Simons et al., 2016). Similar to captopril, rising VEGFA levels led to an increase in vessel density and the number of tip cells (Figure 3E, Figure S2H-N), and shifted the S>R transition from ~230*μ*m to ~260*μ*m (Figure 3F). VEGFA blockage with sFLT1 reduced vascular density, decreased number of tip cells, and abolished the S>R transition (Figure 3G,H, Figure S2H-N), similar to angiotensin-II, yet with a stronger effect. Analysis of individual retinas confirms statistically significant shifts in the S>R transition zone in all conditions, when compared to control retinas (Figure 3I). Vessel morphometrics analyses (independent of cell polarity) confirmed that captopril and VEGFA led to an enrichment in class 1 (sprouting) bins, whereas angiotensin-II or sFLT1 promoted class 2 (remodeling) bins, when compared to control retinas (Figure 3J,K, Figure S3A). Altogether, these results demonstrate that the cellular S>R transition is established by an interaction between blood flow (shear stress) and chemokine (VEGFA) gradients, which compete to define EC collective behavior.

**Figure 3.**
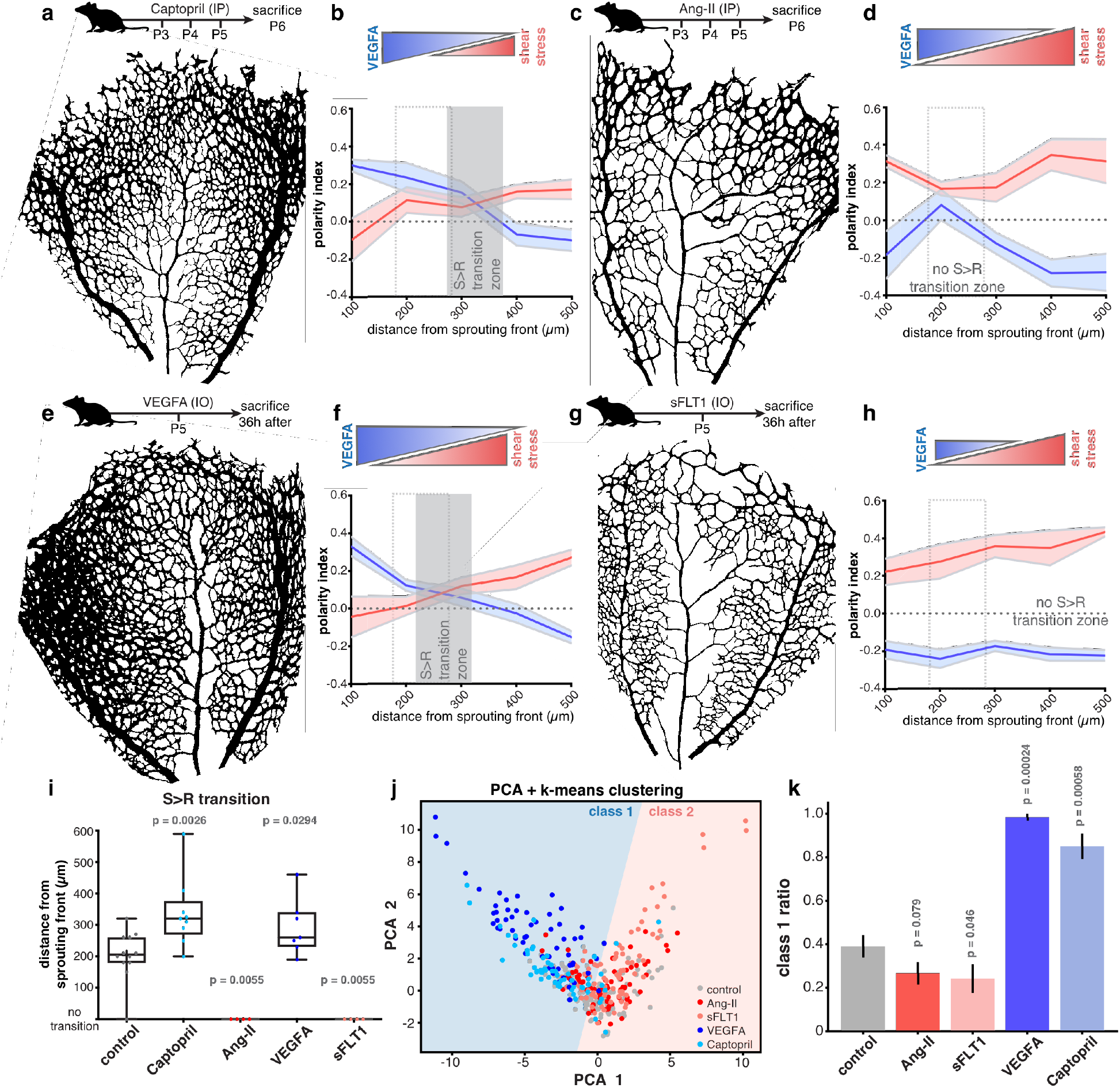
VEGFA and flow pattern govern the S>R transition. **A,C,E,G**, Top: scheme of designated compound injection and collection of samples; Bottom: representative image of the vascular network following designated compound treatment. **B,D,F,H**, Top: proposed strength of each morphogen based on designated compound treatment. Bottom: distribution of K-(blue) and F-(red) indexes within the 500*μ*m from the SF. Solid line represents mean, and light area represents SEM. Dashed grey line represents random polarity. Grey rectangle defines each compound-treated S>R transition. Dashed grey bounding rectangle shows control S>R transition (defined in Figure 2D). **i**, Box plot for the cellular S>R transition in each retina of control, captopril, angiotensin-II, VEGFA and sFLT1 retinas. N=11 control; N=9 captopril; N=4 angiotensin II; N=7 VEGFA; N=4 sFLT1. P-values from Mann-Whitney test between control and the corresponding group. **J**, K-means clustering analysis of vessel morphometrics for designated conditions projected into the binary PCA clustering space defined in Figure 1C. Each dot represents one bin for each retina. **K**, Ratio of class 1 bins over total bins for designated conditions. N= 14 control; N=9 captopril; N=8 angiotensin II; N=7 VEGFA; N=9 sFLT1. P-values of a two-tailed Mann-Whitney test between control and the corresponding group.

### Mechanotransduction at focal adhesions governs shear stress-induced polarity

Numerous reports proposed that EC polarize upstream or downstream in response to flow via different mechanisms (McCue et al., 2006; Tanaka et al., 2021; Tzima et al., 2003) but, the mechanism regulating flow-migration coupling remains unclear. Several shear stress sensors have been identified in ECs, including piezo1, plexinD1, focal adhesions (FAs), the VEGFR2/PECAM1/VE-cadherin complex, or caveolae (Tanaka et al., 2021; Tzima et al., 2005; Xanthis et al., 2019). To clarify how shear stress promotes EC polarity, we establish *in vitro* flow conditions in which ECs polarize against the flow direction, measured by the polarity index (PI) (Carvalho et al., 2019), in a force-dependent manner (Figure S4A,B), as seen *in vivo*. We used 2.0 Pa for 4h as our standard to induce robust collective polarization of EC monolayers. Flow-dependent EC polarization was associated with significant changes in FAs, as previously reported (Jalali et al., 2001; Li et al., 1997). Exposure to shear stress increased the number of FAs per cell, increased their mean length (Figure 4A,B, Figure S4C,D), increased the colocalization of vinculin with integrin alpha 5 (ITGA5) (Figure S4E-G) and of phosphorylated paxillin with vinculin (Figure S5A,B), and increased the phosphorylation of paxillin, vinculin, FAK, and AKT (Figure S5C-E). Remarkably, shear stress induced a bias in the alignment of both FA morphology and the distribution of vinculin-activated integrin beta 1 (aITGB1) colocalization in the direction of flow (Figure 4C,D). Next, we tested the involvement of FAs in EC polarity response. First, we knocked down (KD) either ITGB1, ITGA5, or talin 1, as means to decrease FAs. However, when applying flow, KD cells readily detached, and measurements were not possible. To circumvent this technical issue, we used RGDS, a peptide that binds to integrin RGD-binding motifs, and blocks integrin-mediated adhesion (Kapp et al., 2017). RGDS-treatment impaired FA formation (Figure S6A,B) and led to a significant decrease in EC polarization against the flow direction (Figure 4E). Cilengitide, a RGDS-mimic (Kapp et al., 2017), led to a similar effect (Figure 4E). Thus, FAs are necessary for flow-induced EC polarity. FAs regulate translation-dependent and translation-independent cellular responses. Given that inhibition of transcription, using triptolide, or translation, using puromycin, did not affect flow-mediated polarization (Figure S6C,D), thus polarity response is translation-independent. FAs are mechanoresponsive, sensing the rigidity, molecular composition and conformation of the extracellular matrix (ECM) (Kechagia et al., 2019). To test if mechanotransduction regulates shear stress-induced polarity responses, we manipulated substrate stiffness. Flow-stimulated HUVECs seeded on 3 or 18 kPa soft polydimethylsiloxane (PDMS) had bigger FAs than in static conditions (Figure S7A). Interestingly, shear stress-stimulated ECs at 3kPa showed a tendency for smaller FA and a lower PI compared to those at 18kPa (Figure S7A,B). The relationship between stiffness and flow-induced PI was even more evident in stiffer substrates. Stiffer hard PDMS (1MPa) had higher PI compared to softer hard PDMS (280kPa), a phenomenon that was associated positively with the length of FAs in the different conditions (Figure S7C,D). These results suggest that the ability of ECs to polarize against the flow direction is dependent on the strength of adhesion to the ECM. To confirm this hypothesis, we measured FA-mediated tension through traction force microscopy (TFM) (Butler et al., 2002). As expected, direct application of flow on the substrate without cells does not generate measurable tractions (Figure S7E,F). TFM revealed that flow-stimulated ECs exerted significantly higher levels of traction forces on the substrate, when compared to static conditions (Figure 4F,G). However, the amount of tension per vinculin molecule, estimated by FRET efficiency, was similar under flow or static conditions (Figure S7G,H). FA maturation and traction forces are mediated by myosin-II-dependent contractility (Chrzanowska-Wodnicka and Burridge, 1996). Inhibition of ROCK (Y-27632) or myosin ATPase activity (blebbistatin - BBS) led to a significant decrease in the number and size of FAs (Figure 4H, Figure S8A,B), and in the colocalization of vinculin with ITGA5 (Figure S8C), which was associated with a decrease in traction forces, when compared to control conditions (compare Figure 4G and Figure S8E). These effects correlated with a significant impairment in flow-induced EC polarization under actomyosin inhibition (Figure 4I). Thus, our data suggest that shear stress stimulates FA assembly and enhances traction forces, which are required for flow-induced EC polarization.

**Figure 4.**
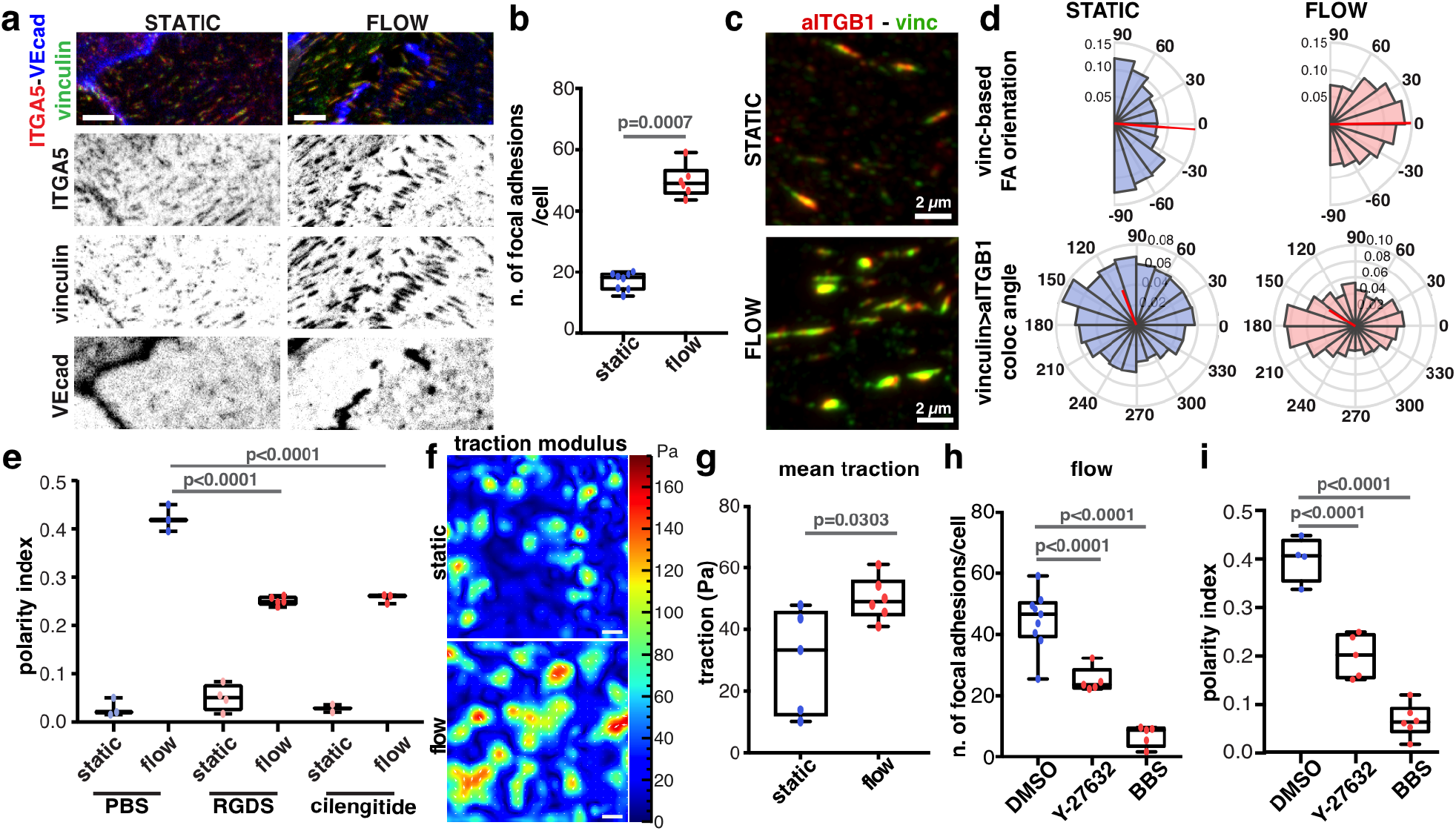
FA-mediated traction forces drive flow-induced polarity. **A**, Representative image of AJs (VE-cadherin – VEcad, blue) and FAs (vinculin, green; and integrin alpha 5 – ITGA5, red) in static or flow-stimulated HUVEC monolayers (high-magnification from Figure S4C). Scale bar: 2μm. **B**, Box plot for the number of FAs in static or flow conditions. N=8 (static) and N=6 (flow). p-value from Mann-Whitney test. **C**, Representative super-resolution image of co-localization between vinculin (green), and activated integrin beta 1 – aITGB1 (red) in static or flow-stimulated HUVEC monolayers. Scale bar: 2μm. **D**, Top: Angular histograms for the distribution of vinculin FA orientation in static and flow conditions. Bottom: mean angle of orientation of the vector from vinculin to aITGB1 centroids in relation to right-to-left slide axis in static and flow conditions. Flow direction is right-to-left. N=20 images, 4 independent experiments. **E**, Box plot for polarity index in static or flow-stimulated HUVEC monolayers treated with PBS (N=3), RGDS (N=5) or cilengitide (N=3); p-values from one-way ANOVA with Sidak test. **F,G**, Mean traction maps (F) and box plot of mean traction forces (G) exerted by static or flow-stimulated HUVEC monolayers. N=4 static; N=5 flow; p-values from Mann-Whitney test. **H**, Box plot for number of focal adhesions in flow-stimulated HUVEC monolayers treated with DMSO (N=9), Y-27632 (N=5) or BBS (blebbistatin, N=5). p-values from Mann-Whitney test. **i**, Box plot of polarity index in static or flow-stimulated HUVEC monolayers treated with DMSO (N=4), Y-27632 (N=5) or BBS (blebbistatin, N=6). p-values from one-way ANOVA with Sidak test.

### A shift in tension distribution regulates the competition between chemokines and shear stress to establish EC polarity patterns

Our data, together with previous reports, suggest that collective EC polarity is regulated by chemokines through tension at AJs (Carvalho et al., 2019; Hayer et al., 2016; Huveneers et al., 2012), whilst shear stress acts through tension at FAs (Figure 5A). To experimentally test this hypothesis, we established an *in vitro* assay in which wounded HUVEC monolayers are exposed or not to flow (Figure 5B). The wound in the monolayer generates a free edge that stimulates collective migration similarly to chemokines (Carvalho et al., 2019; Friedl and Mayor, 2017; Vitorino and Meyer, 2008). We used the scratch-wound assay rather than VEGFA gradients because counter-gradients of soluble factors are difficult to establish under high flow rate (20mL/min) conditions. KD of VEGFR2 inhibited free edge induced polarity whilst not affecting flow-induced polarity (Figure S9A,B), confirming that scratch-wound assay is a suitable surrogate to investigate chemokine-induced polarity. In this competition assay, we determined the PI of ECs in ~200*μ*m wide regions on the left (convergent), on the right (divergent), and in regions >2mm away (non-interactive) from the wound (Figure 5B). Under static conditions, ECs were unpolarized (PI= ~ 0.0) in non-interactive regions, whilst convergent and divergent regions had mirrored responses (PI of 0.23 and −0.24, respectively) (Figure 5C). Shear stress (2.0Pa) induced robust EC polarity in all the regions (Figure 5C), suggesting that shear stress is a stronger polarity cue than the free edge. Yet, instead of a homogenous flow response, we observed a statistically significant lower PI (0.33) in divergent compared to non-interactive regions (0.39), which was also significantly lower than in the convergent region (0.46) (Figure 5C). These results suggest that each polarity cue triggers independent pathways, which can lead to competitive or cooperative effects on cell polarity depending on the directionality of each polarity cue.

**Figure 5.**
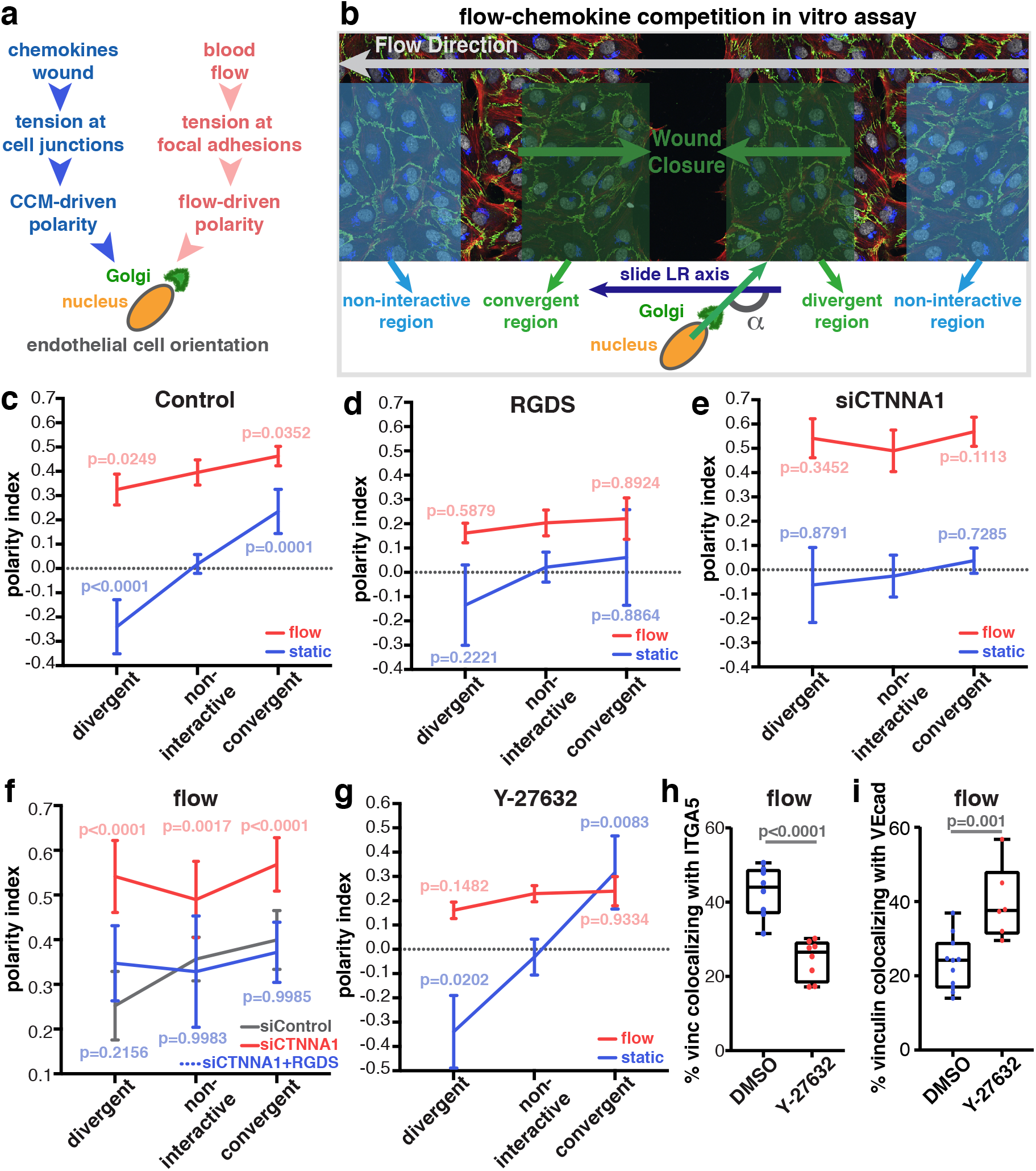
Chemokines and blood flow compete to establish the main polarity axis of ECs. **A**, Schematic in chemokine- and flow-induced collective polarity of ECs. **B**, Layout of the flow-chemokine competition *in vitro* assay, depicting the relationship between each wound side and the flow direction. EC polarity angle (α) is measured in relation to the right-to-left (RL) axis of the slide, in the same direction of flow. **C-E**, PI for designated regions in static (blue) or flow (red) in control (C), RGDS (D) and siCTNNA1 (E) HUVECs. N=4 Control; N=5 RGDS; N=7 siCTNNA1. p-values for multiple comparisons with non-interactive region one-way ANOVA with Sidak test. **F**, PI for designated regions in static or flow conditions in siControl (red) or siCTNNA1 HUVECs treated with PBS (grey) or RGDS (blue). N=4 per condition; red p-values (siCTNNA1) and blue p-values (siCtnna1 + RGDS) for multiple comparisons with siControl one-way ANOVA with Sidak test. **G**, PI for designated regions in static (blue) or flow (red) conditions in HUVECs treated with DMSO or Y-27632. N=5 per condition. p-values correspond to multiple comparisons with non-interactive region one-way ANOVA with Sidak test. **H**, Box plot for percentage for vinculin colocalizing with in ITGA5 in flow-stimulated DMSO- or Y-27632-treated HUVEC monolayers. N=8 per condition; p-values from Mann-Whitney test. **I**, Box plot of percentage of vinculin colocalizing with VE-cadherin (VEcad) in flow-stimulated HUVEC monolayers treated with DMSO (N=11) or Y-27632 (N=6). P-values from Mann-Whitney test.

Next, we perturbed FAs and AJs, and tested their effects in the competition assay. Inhibition of FA formation, through RGDS treatment, disrupted free edge- and flow-induced EC polarity, when compared to control cells (compare Figure 5C to Figure 5D). SiRNA-mediated KD of alpha-catenin (siCTNNA1) (Figure S9C), a crucial protein of AJs (Bazellières et al., 2015), led to the expected inhibition of free edge-induced polarity in static conditions (Figure 5E), but, surprisingly, also led to a significant sensitization to shear stress when compared to siControl cells (Figure 5E, Figure S9D). This effect was further confirmed by KD of VE-cadherin (siCDH5) (Figure S9E). siRNA-mediated depletion of PECAM1 or VEGFR2 demonstrate that AJs regulate flow-induced EC polarization independently of the well-known VE-cadherin-PECAM1-VEGFR2 mechanosensory complex (Tzima et al., 2005) (Figure S9B,F). Together, these results suggest that AJs are positive regulators of free edge-induced polarization and negative regulators of flow-induced polarization. Mechanical feedback between AJ- and FA-based adhesion has been previously documented (Hur et al., 2012; Liu et al., 2010; Mertz et al., 2013; Tambe et al., 2011). Thus, we hypothesized that disruption of AJs could stimulate flow-dependent polarization by promoting FA-dependent mechanotransduction. Supporting this possibility, we observed that siCTNNA1 cells have more and larger FAs than control cells in static and flow conditions (Figure S10A-C). In agreement, inhibition of FAs in siCTNNA1 cells with RGDS led to a significant reduction of flow-mediated polarity, to levels indistinguishable from siControl cells (Figure 5F). Additionally, shear stress increased significantly vinculin co-localization with FAs (Figure S10D), a protein known to be recruited to both FAs and AJs in a force-dependent manner (Bays and DeMali, 2017), which was mirrored by a significant decrease in AJs (Figure S10E). This suggests a shift in the distribution of mechanical stresses upon flow exposure. In agreement, siCTNNA1 cells display an increase in co-localization of vinculin with ITGA5 in static conditions, whilst no significant changes were observed under flow (Figure S10D). Remarkably, monolayer stress microscopy (Tambe et al., 2011) demonstrated that intercellular tension was increased upon flow exposure, in accordance with a previous report (Hur et al., 2012) (Figure S11A,B). Taken together, these observations suggest that flow promotes tension both at FAs and AJs, but that flow-dependent tension at FAs is a stronger polarity stimulus than tension at AJs, a response that is further enhanced when AJs are disrupted. In accordance, inhibition of actomyosin contractility by Y-27632 negatively regulated flow-induced polarity in divergent, non-interactive and convergent regions (Figure 5G). Yet, Y-27632 did not significantly alter free edge-induced polarity (p=0.1208 in divergent; p=0.7594 in non-interactive; p=0.2168 in convergent, compare Figure 5C and Figure 5G). The effect of Y-27632 on PIs correlated with a decrease in the colocalization between vinculin and integrin alpha5 (Figure 5H), and an increase in the colocalization of vinculin with VE-cadherin under flow conditions (Figure 5I). Taken together, these observations strongly suggest that shear stress increases ROCK-dependent FA-derived polarity cues, and reduces the ability of cells to coordinate free edge collective behaviors that rely on tension at AJs.

Next, we sought to integrate these *in vitro* observations into the physiological context of angiogenesis by manipulating tension at FAs and AJs, and to measure their effects on the S>R transition. Inhibition of integrin-mediated binding to the ECM using RGDS *in vivo* led to highly variable transition zones, associated with an impairment in both K- and F-indexes (Figure 6A, Figure S12A,B), likely because RGDS treatment affects both chemokine-induced polarity and flow-induced polarity (Figure 5D). Inhibition of ROCK-induced actomyosin contractility significantly delayed the S>R transition zone (Figure 6B, Figure S12C,D), similarly to *in vitro* (Figure 5G). These results were further confirmed using an endothelial-specific deletion of Myh9, coding for the heavy chain of the myosin-IIA (Figure 6C, Figure S12E,F). Conversely, disruption of AJ-mediated tension by endothelial-specific deletion of Ctnna1 promoted flow-dependent polarity whereas inhibiting chemokine-induced polarity (Figure 6D, Figure S12G-I). Taken together, we propose a theoretical model for the establishment of the S>R transition zone based on the gradual increase of flow-mediated tension at FAs, which promotes a shift from AJ-mediated polarity towards FA-mediated polarity (Figure 6E).

**Figure 6.**
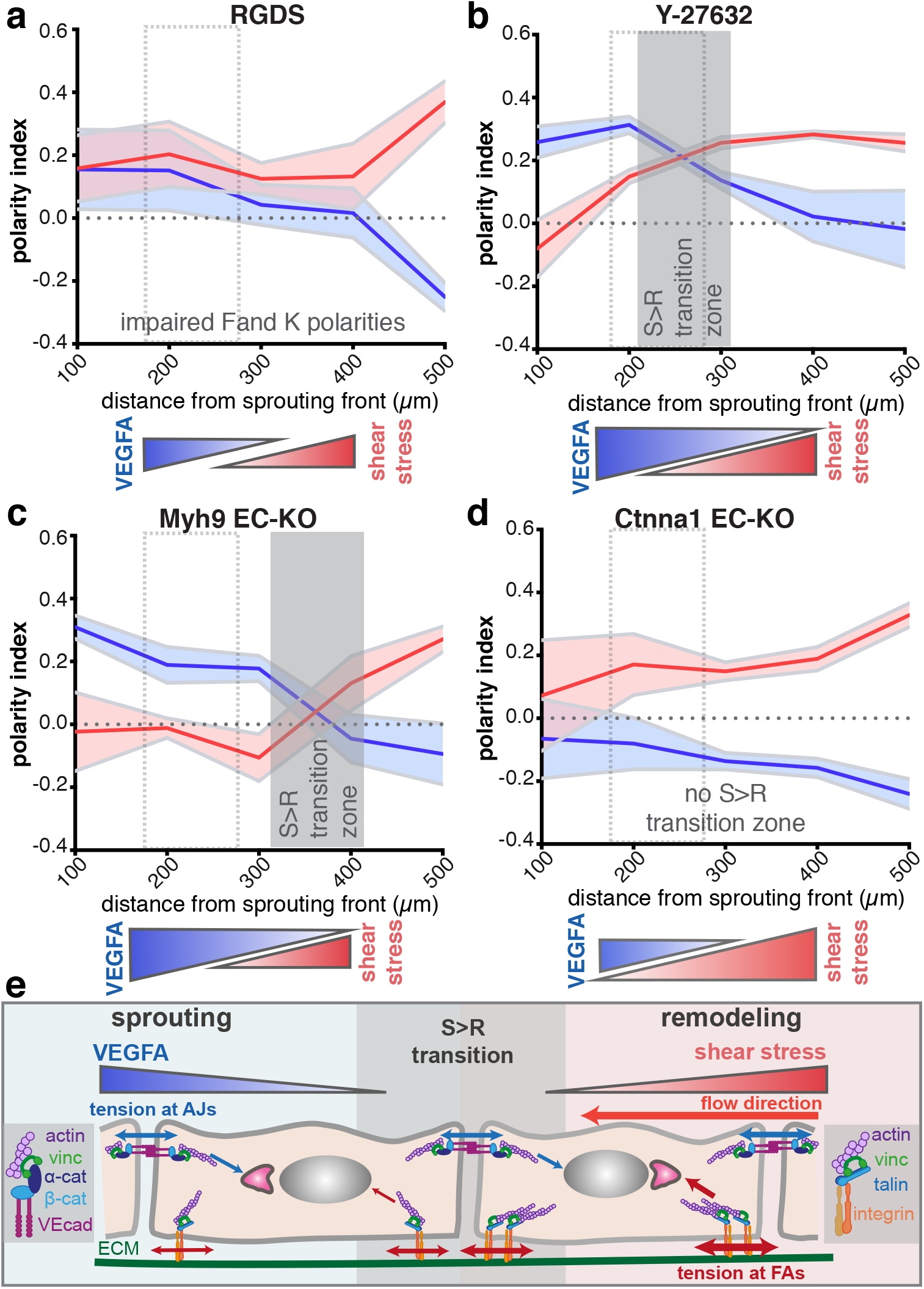
Manipulation of AJs or FAs alters the S>R transition zone. **A-D**, Top: distribution of K-(blue) and F-(red) indexes within the 500*μ*m from the SF. Bottom: proposed strength of each morphogen based on designated drug treatment or KO animal. Solid line represents mean, and light area represents SEM. Dashed grey line represents random polarity. Grey rectangle defines each drug-treated or KO animal S>R transition. Dashed grey bounding rectangle shows control S>R transition (defined in Figure 2D). N=5 RGDS; N=5 Y-27632; N=5 Myh9 EC-KO; N=4 Ctnna1 EC-KO; arterial regions. **E**, Mechanical regulation of the S>R transition.

## Discussion

Here, we propose that the competitive regulation of cell polarity leads to a shift in cell-level migratory behavior (toward VEGFA or against blood flow), which translates into a tissue-level morphogenetic event, the S>R transition. This simple principle enables local vascular adaptation that could underlie the generation of unique vascular patterns. As VEGFA levels and flow patterns differ in each organ during development and in pathological conditions, our model could provide a biological explanation for organotypic vasculature and why these patterns become disrupted in disease (Augustin and Koh, 2017).

We have identified the S>R transition based on cell polarity, which could be explained largely by a biomechanical perspective. Nevertheless, shear stress levels and pressure differences does also influence gene expression, cell identity, and sprouting activity of ECs (Chouinard-Pelletier et al., 2013; Ghaffari et al., 2015; Song and Munn, 2011). Moreover, besides VEGFA, several other signaling molecules further regulate vascular biology, such as BMP9/10, S1P and ANG2 (Potente and Makinen, 2017). Our S>R transition model could provide powerful tool to explore the interconnections between mechanical and biochemical signaling and refine our current. In this context, one interesting biological process is arteriogenesis, in which pre-arterial ECs differentiate at the sprouting front and migrate upstream to contribute to artery formation (Hasan et al., 2017; Pitulescu et al., 2017; Su et al., 2018). Determining how and why pre-arterial ECs are more effective at flow-migration coupling than neighboring ECs can have important implications for vascular morphogenesis. NOTCH, CXCR4, BMP signaling have been linked to arteriogenesis and/or to modulate EC shear stress responses (Baeyens et al., 2015; Cartier and Hla, 2019; Mack et al., 2017; Tanaka et al., 2021), and thus are prime candidates to fine-tune the S>R transition at the single cell level. Moreover, computational modeling of adaptation and maladaptation showed that well organized vascular networks rely on tight feedback between vessel caliber control and hemodynamics (Alberding and Secomb, 2021; Pries et al., 2009), and transfer of information from distal to proximal vessels in arteries (Pries et al., 2010). How these flows of information relate to EC polarity and the S>R transition remains to be explored.

Investigating the competitive and synergistic interactions driving cell polarity and the interplay between biochemical and physical crosstalk could offer opportunities to modulate the S>R transition and regulate vascular morphogenesis in health and disease.

## Acknowledgements

We thank the VML lab and Fondation Leducq ATTRACT members for helpful discussions. We thank Isabel Flores for helping with statistical analysis. Supercomputing time on the ARCHER UK National Supercomputing Service was provided by the “UK Consortium on Mesoscale Engineering Sciences (UKCOMES)”

## Funding

European Research Council (679368): CAF.

European Research Council (883739): XT.

Fundação para a Ciência e Tecnologia (PTDC/MED-PAT/31639/2017; PTDC/BIA-CEL/32180/2017; CEECIND/04251/2017): CAF.

Fondation LeDucq (17CVD03): CAF, MOB.

European Commission (801423): CAF, MOB.

European Commission (731957): XT, PRC.

EPSRC (EP/T008806/1; EP/R029598/1): MOB.

Spanish Ministry of Science and Innovation (PID2019-110298GB-I00): PRC.

Spanish Ministry of Science and Innovation (PGC2018-099645-B-I00): XT.

Generalitat de Catalunya (2017-SGR-1602): XT, PRC.

La Caixa Foundation (LCF/PR/HR20/52400004): XT, PRC.

Fundació la Marató de TV3 (201903-30-31-32): XT.

EMBO (ALTF 2-2018): LMF.

EU MSCA (842498): MO

## Author Contributions

Conceptualization: PB, MDC, XT, PRC, MOB, CAF.

Methodology: PB, MDC, CGF, GZ, LMF, MO, MG, PRC, MOB, and CAF

Experimentation and data acquisition: PB, MDC, CGF, GZ, AAP, AP, DR, YG, MO, DB, IF, LM, IM, YC, CAF

Data analysis: PB, MDC, CGF, LMF, MGG, YG, LM, XT, PRC, AE, MOB, CAF

Project administration: CAF

Writing – original draft: CAF

Writing – review & editing: PB, CGF, LMF, MO, MG, AP, IF, XT, PRC, AE, MOB, and CAF

## Data and materials availability

All data are available in the main text or the supplementary materials.

## Code availability

In this manuscript, we used mathematical algorithm. PolNet was published previously (ref). Code for Vessel Morphometrics can be access upon request to Miguel Bernabeu. Code for Traction Force Microscopy and Monolayer Stress Microscopy can be access upon request to Xavier Trepat.

## References

Alberding, J.P., and Secomb, T.W. (2021). Simulation of angiogenesis in three dimensions: Application to cerebral cortex. PLoS computational biology 17, e1009164.

Augustin, H.G., and Koh, G.Y. (2017). Organotypic vasculature: From descriptive heterogeneity to functional pathophysiology. Science 357.

Baeyens, N., Nicoli, S., Coon, B.G., Ross, T.D., Van den Dries, K., Han, J., Lauridsen, H.M., Mejean, C.O., Eichmann, A., Thomas, J.L., et al. (2015). Vascular remodeling is governed by a VEGFR3-dependent fluid shear stress set point. Elife 4.

Bays, J.L., and DeMali, K.A. (2017). Vinculin in cell-cell and cell-matrix adhesions. Cell Mol Life Sci 74, 2999–3009.

Bazellières, E., Conte, V., Elosegui-Artola, A., Serra-Picamal, X., Bintanel-Morcillo, M., Roca-Cusachs, P., Muñoz, J.J., Sales-Pardo, M., Guimerà, R., and Trepat, X. (2015). Control of cell-cell forces and collective cell dynamics by the intercellular adhesome. Nat Cell Biol 17, 409–420.

Bernabeu, M.O., Jones, M.L., Nash, R.W., Pezzarossa, A., Coveney, P.V., Gerhardt, H., and Franco, C.A. (2018). PolNet: A Tool to Quantify Network-Level Cell Polarity and Blood Flow in Vascular Remodeling. Biophys J 114, 2052–2058.

Butler, J.P., Tolić-Nørrelykke, I.M., Fabry, B., and Fredberg, J.J. (2002). Traction fields, moments, and strain energy that cells exert on their surroundings. American journal of physiology Cell physiology 282, C595–605.

Cao, J., Ehling, M., März, S., Seebach, J., Tarbashevich, K., Sixta, T., Pitulescu, M.E., Werner, A.C., Flach, B., Montanez, E., et al. (2017). Polarized actin and VE-cadherin dynamics regulate junctional remodelling and cell migration during sprouting angiogenesis. Nat Commun 8, 2210.

Cartier, A., and Hla, T. (2019). Sphingosine 1-phosphate: Lipid signaling in pathology and therapy. Science 366.

Carvalho, J.R., Fortunato, I.C., Fonseca, C.G., Pezzarossa, A., Barbacena, P., Dominguez-Cejudo, M.A., Vasconcelos, F.F., Santos, N.C., Carvalho, F.A., and Franco, C.A. (2019). Non-canonical Wnt signaling regulates junctional mechanocoupling during angiogenic collective cell migration. Elife 8.

Chouinard-Pelletier, G., Jahnsen, E.D., and Jones, E.A. (2013). Increased shear stress inhibits angiogenesis in veins and not arteries during vascular development. Angiogenesis 16, 71–83.

Chrzanowska-Wodnicka, M., and Burridge, K. (1996). Rho-stimulated contractility drives the formation of stress fibers and focal adhesions. J Cell Biol 133, 1403–1415.

Fonseca, C.G., Barbacena, P., and Franco, C.A. (2020). Endothelial cells on the move: dynamics in vascular morphogenesis and disease. Vascular biology (Bristol, England) 2, H29–h43.

Franco, C.A., Jones, M.L., Bernabeu, M.O., Geudens, I., Mathivet, T., Rosa, A., Lopes, F.M., Lima, A.P., Ragab, A., Collins, R.T., et al. (2015). Dynamic endothelial cell rearrangements drive developmental vessel regression. PLoS Biol 13, e1002125.

Friedl, P., and Mayor, R. (2017). Tuning Collective Cell Migration by Cell-Cell Junction Regulation. Cold Spring Harb Perspect Biol 9.

Ghaffari, S., Leask, R.L., and Jones, E.A. (2015). Flow dynamics control the location of sprouting and direct elongation during developmental angiogenesis. Development 142, 4151–4157.

Hartung, D., Olsen, M.A., Xu, H., Nguyen, H.T., and Busch, C. (2012). Comprehensive analysis of spectral minutiae for vein pattern recognition. In IET Biometrics (Institution of Engineering and Technology), pp. 25–36.

Hasan, S.S., Tsaryk, R., Lange, M., Wisniewski, L., Moore, J.C., Lawson, N.D., Wojciechowska, K., Schnittler, H., and Siekmann, A.F. (2017). Endothelial Notch signalling limits angiogenesis via control of artery formation. Nat Cell Biol 19, 928–940.

Hayer, A., Shao, L., Chung, M., Joubert, L.M., Yang, H.W., Tsai, F.C., Bisaria, A., Betzig, E., and Meyer, T. (2016). Engulfed cadherin fingers are polarized junctional structures between collectively migrating endothelial cells. Nat Cell Biol 18, 1311–1323.

Hur, S.S., del Álamo, J.C., Park, J.S., Li, Y.S., Nguyen, H.A., Teng, D., Wang, K.C., Flores, L., Alonso-Latorre, B., Lasheras, J.C., et al. (2012). Roles of cell confluency and fluid shear in 3-dimensional intracellular forces in endothelial cells. Proc Natl Acad Sci U S A 109, 11110–11115.

Huveneers, S., Oldenburg, J., Spanjaard, E., van der Krogt, G., Grigoriev, I., Akhmanova, A., Rehmann, H., and de Rooij, J. (2012). Vinculin associates with endothelial VE-cadherin junctions to control force-dependent remodeling. J Cell Biol 196, 641–652.

Jalali, S., del Pozo, M.A., Chen, K., Miao, H., Li, Y., Schwartz, M.A., Shyy, J.Y., and Chien, S. (2001). Integrin-mediated mechanotransduction requires its dynamic interaction with specific extracellular matrix (ECM) ligands. Proc Natl Acad Sci U S A 98, 1042–1046.

Kapp, T.G., Rechenmacher, F., Neubauer, S., Maltsev, O.V., Cavalcanti-Adam, E.A., Zarka, R., Reuning, U., Notni, J., Wester, H.J., Mas-Moruno, C., et al. (2017). A Comprehensive Evaluation of the Activity and Selectivity Profile of Ligands for RGD-binding Integrins. Sci Rep 7, 39805.

Kechagia, J.Z., Ivaska, J., and Roca-Cusachs, P. (2019). Integrins as biomechanical sensors of the microenvironment. Nat Rev Mol Cell Biol 20, 457–473.

Korn, C., and Augustin, H.G. (2015). Mechanisms of Vessel Pruning and Regression. Dev Cell 34, 5–17.

Kwon, H.B., Wang, S., Helker, C.S., Rasouli, S.J., Maischein, H.M., Offermanns, S., Herzog, W., and Stainier, D.Y. (2016). In vivo modulation of endothelial polarization by Apelin receptor signalling. Nat Commun 7, 11805.

Li, S., Kim, M., Hu, Y.L., Jalali, S., Schlaepfer, D.D., Hunter, T., Chien, S., and Shyy, J.Y. (1997). Fluid shear stress activation of focal adhesion kinase. Linking to mitogen-activated protein kinases. J Biol Chem 272, 30455–30462.

Liu, Z., Tan, J.L., Cohen, D.M., Yang, M.T., Sniadecki, N.J., Ruiz, S.A., Nelson, C.M., and Chen, C.S. (2010). Mechanical tugging force regulates the size of cell-cell junctions. Proc Natl Acad Sci U S A 107, 9944–9949.

Mack, J.J., Mosqueiro, T.S., Archer, B.J., Jones, W.M., Sunshine, H., Faas, G.C., Briot, A., Aragón, R.L., Su, T., Romay, M.C., et al. (2017). NOTCH1 is a mechanosensor in adult arteries. Nat Commun 8, 1620.

McCue, S., Dajnowiec, D., Xu, F., Zhang, M., Jackson, M.R., and Langille, B.L. (2006). Shear stress regulates forward and reverse planar cell polarity of vascular endothelium in vivo and in vitro. Circ Res 98, 939–946.

Mertz, A.F., Che, Y., Banerjee, S., Goldstein, J.M., Rosowski, K.A., Revilla, S.F., Niessen, C.M., Marchetti, M.C., Dufresne, E.R., and Horsley, V. (2013). Cadherin-based intercellular adhesions organize epithelial cell-matrix traction forces. Proc Natl Acad Sci U S A 110, 842–847.

Nehme, A., Zouein, F.A., Zayeri, Z.D., and Zibara, K. (2019). An Update on the Tissue Renin Angiotensin System and Its Role in Physiology and Pathology. Journal of cardiovascular development and disease 6.

Park, H., Furtado, J., Poulet, M., Chung, M., Yun, S., Lee, S., Sessa, W.C., Franco, C.A., Schwartz, M.A., and Eichmann, A. (2021). Defective Flow-Migration Coupling Causes Arteriovenous Malformations in Hereditary Hemorrhagic Telangiectasia. Circulation.

Pitulescu, M.E., Schmidt, I., Giaimo, B.D., Antoine, T., Berkenfeld, F., Ferrante, F., Park, H., Ehling, M., Biljes, D., Rocha, S.F., et al. (2017). Dll4 and Notch signalling couples sprouting angiogenesis and artery formation. Nat Cell Biol 19, 915–927.

Potente, M., and Makinen, T. (2017). Vascular heterogeneity and specialization in development and disease. Nat Rev Mol Cell Biol 18, 477–494.

Pries, A.R., Cornelissen, A.J., Sloot, A.A., Hinkeldey, M., Dreher, M.R., Höpfner, M., Dewhirst, M.W., and Secomb, T.W. (2009). Structural adaptation and heterogeneity of normal and tumor microvascular networks. PLoS computational biology 5, e1000394.

Pries, A.R., Höpfner, M., le Noble, F., Dewhirst, M.W., and Secomb, T.W. (2010). The shunt problem: control of functional shunting in normal and tumour vasculature. Nature reviews Cancer 10, 587–593.

Simons, M., Gordon, E., and Claesson-Welsh, L. (2016). Mechanisms and regulation of endothelial VEGF receptor signalling. Nat Rev Mol Cell Biol 17, 611–625.

Song, J.W., and Munn, L.L. (2011). Fluid forces control endothelial sprouting. Proc Natl Acad Sci U S A 108, 15342–15347.

Su, T., Stanley, G., Sinha, R., D’Amato, G., Das, S., Rhee, S., Chang, A.H., Poduri, A., Raftrey, B., Dinh, T.T., et al. (2018). Single-cell analysis of early progenitor cells that build coronary arteries. Nature 559, 356–362.

Tambe, D.T., Hardin, C.C., Angelini, T.E., Rajendran, K., Park, C.Y., Serra-Picamal, X., Zhou, E.H., Zaman, M.H., Butler, J.P., Weitz, D.A., et al. (2011). Collective cell guidance by cooperative intercellular forces. Nature materials 10, 469–475.

Tanaka, K., Joshi, D., Timalsina, S., and Schwartz, M.A. (2021). Early events in endothelial flow sensing. Cytoskeleton (Hoboken, NJ).

Tzima, E., del Pozo, M.A., Shattil, S.J., Chien, S., and Schwartz, M.A. (2001). Activation of integrins in endothelial cells by fluid shear stress mediates Rho-dependent cytoskeletal alignment. Embo j 20, 4639–4647.

Tzima, E., Irani-Tehrani, M., Kiosses, W.B., Dejana, E., Schultz, D.A., Engelhardt, B., Cao, G., DeLisser, H., and Schwartz, M.A. (2005). A mechanosensory complex that mediates the endothelial cell response to fluid shear stress. Nature 437, 426–431.

Tzima, E., Kiosses, W.B., del Pozo, M.A., and Schwartz, M.A. (2003). Localized cdc42 activation, detected using a novel assay, mediates microtubule organizing center positioning in endothelial cells in response to fluid shear stress. J Biol Chem 278, 31020–31023.

Vitorino, P., and Meyer, T. (2008). Modular control of endothelial sheet migration. Genes Dev 22, 3268–3281.

Xanthis, I., Souilhol, C., Serbanovic-Canic, J., Roddie, H., Kalli, A.C., Fragiadaki, M., Wong, R., Shah, D.R., Askari, J.A., Canham, L., et al. (2019). β1 integrin is a sensor of blood flow direction. J Cell Sci 132.

