## Supplemental Methods and Figures for "Competition for endothelial cell polarity drives vascular morphogenesis"

### Supplementary Information

#### Mice and Treatments

In this study, we used the following mouse strains: Myh9 floxed[1]; Ctnna1 floxed[2]; PdgfbiCreERT2[3]; Cdh5CreERT2[4]; and WT C57BL/6. C57BL/6 pups were used for modulation of flow, VEGF availability or drug treatment. The phenotype of Myh9 endothelial-specific deletion was characterized previously[5], whilst endothelial-specific deletion of Ctnna1 will be characterized in a future publication. For increasing blood flow, Angiotensin II (Sigma-Aldrich, A9525) was injected intraperitoneally (IP) 10µL/g (10 mg/mL stock solution) daily at postnatal day 3 (P3), P4 and P5 and pups were collected at P6. On the opposite, to reduce blood flow, Captopril (Sigma-Aldrich, C4042) was injected IP 15µL/g (3.3mg/mL stock solution) daily at P3, P4 and P5 and pups were collected at P6. In both experiments, control mice were injected using PBS alone. For intraocular administration of reagents, P4-P5 pups were anesthetized and intravitreal injections were performed under a stereomicroscope using a 10µl Hamilton syringe equipped with a 33-gauge needle. Approximately 0.5µl of sterile solution or PBS was injected per eye, while the contralateral eye remained uninjected. The following substances were used: recombinant VEGFA (493-VE-050; R&D Systems, 3µg/µl) or recombinant VEGFR1/Flt-1 Fc chimera protein (sFLT1; 471-F1-100; R&D Systems, 1µg/µl). Mice were sacrificed 36h later. The eyes were removed, fixed in 2% paraformaldehyde (PFA, Sigma-Aldrich, 4412244) in PBS at 4ºC for 5h, rinsed in PBS and processed for IHC.

In the case of the drug experiments, Y-27632 (Merck Millipore, 688001) (10mg/kg) and RGDS (Tocris, 3498) (5mg/kg) were both injected IP at P5 and pups were sacrificed at P6.

In gene deletion experiments, 4-hydroxytamoxifen (Sigma-Aldrich, H6278) was injected IP (20 µg/g) at P1 and P3 for Myh9 floxed mice and at P4 for Ctnna1 floxed, and eyes were collected at P6. As controls, Cre-negative littermates were used in all experiments. Both males and females were used, without distinction.

Mice were maintained at the Instituto de Medicina Molecular (iMM) under standard husbandry conditions and under national regulations. Animal procedures were performed under the DGAV project license 0421/000/000/2016.

#### Immunofluorescence on Mouse Retinas

Retinas were stained as previously described[6]. Briefly, retinas were incubated for 2h, at RT in Claudio's Blocking Buffer (CBB). Next, they were incubated with anti-ICAM2 (1:100, BD Biosciences 553326) and anti-ERG (1:200, Abcam ab92513) primary antibodies in 1:1 PBS and CBB mixture at 4ºC o/n. On the next day, retinas were washed 3 times (30min each) with PBST (PBS with 0.1% Triton X-100, Sigma-Aldrich, T8787) and further incubated with secondary antibodies anti-rat Alexa 555 (Thermo Fisher Scientific, A21434) and anti-rabbit Alexa 488 (Thermo Fisher Scientific, A21206) in 1:1 PBS and CBB mixture o/n at 4ºC. The day after, retinas were washed three times (30min each) with PBST and incubated with $Fab_2$ fragments (Donkey anti-rabbit, Jackson Immuno Research, 711-006-152) for 2h at RT with gentle shaking. Afterwards retinas were fixed with 4% paraformaldehyde (PFA) at RT for 15min and washed three times (15min each) with PBST, followed by 1h incubation with CBB at RT. Next, retinas were incubated with anti-GOLPH4 primary antibody (1:400, Abcam, ab28049) in 1:1 PBS and CBB mixture at 4ºC o/n. The day after, retinas were washed three times (30min each) with PBST and incubated with anti-rabbit Alexa 647 (Thermo Fisher Scientific, A31573) secondary antibody in 1:1 PBS and CBB o/n at 4ºC. Then, retinas were washed three times (30min each) with PBST and flat-mounted on glass slides using Vectashield mounting medium

(Vector Laboratories, H-1000). Images were acquired by tile-scans with multiple Z-positions using a Zeiss Cell Observer SD (Carl Zeiss) equipped with Zen software and with a Plan-Apochromat 40x NA 1.40 oil objective.

**Culture of HUVECs**

Human umbilical vein endothelial cells – HUVECs (Lonza, C2519A) – were cultured following the manufacturer's guidelines, in filter-cap T75 flasks Nunclon Δ surface treatment (VWR international, LLC) with complete medium EGM-2 Bulletkit (Lonza, CC-3162) at 37ºC and 5% $CO_2$ to ensure stable environment for optimal cell growth. All the experiments were conducted with HUVECs between passages 1 and 5. When passaging HUVECs for experiments, cells were washed twice in sterile PBS (137mM NaCl, 2.7mM KCl, 4.3mM $Na_2HPO_4$, 1.47mM $KH_2PO_4$, pH7.4) and then incubated for 5min in TrypLE™ Express Enzyme (1X) (Alfagene, 12605028) at 37ºC, 5% $CO_2$. When 95% of the cells detached, complete medium was added to each flask to inhibit the activity of the TrypLE™ Express Enzyme and the cell suspension was transferred into a falcon tube. Cells were then centrifuged at 700rpm for 5min at RT and the pellet re-suspended in fresh complete medium. HUVECs were then seeded at the desired concentration, depending on the experiments.

**siRNA Transfection**

In order to silence the expression of genes of interest, a set of ON-TARGET human siRNAs against CTNNA1 (Dharmacon, GE Healthcare, J-010505-06), CDH5 (Dharmacon, GE Healthcare, J-003641-07) or untargeting control were used. Briefly, HUVECs were seeded the day before the transfection to reach 60-70% confluence and were then transfected with 25nM of siRNA using the DharmaFECT 1 reagent (Dharmacon, GE Healthcare) following the Dharmacon siRNA Transfection Protocol. 24h after transfection the culture medium was replaced by fresh complete medium and cells were kept under culture conditions up until 72h post-transfection and then processed for further experiments.

**Viral Production and Transduction**

Replication-incompetent lentiviruses were produced by transient transfection of HEK293T with lentiviral expression vector co-transfected with the viral packaging vector Δ8.9 and the viral envelope vector VSVG. Medium was replaced with fresh culture medium 6-8h post transfection. 48h after medium replacement, lentiviral particles were concentrated from supernatant by ultracentrifugation at 112.500g for 1h30 and re-suspended in 0.1% BSA PBS. Seeded HUVECs were transduced with varying concentrations of a lentiviral plasmid containing pRRL-VinculinTS [7]. 24h after viral transduction the culture medium was replaced by fresh complete medium and cells were kept under culture conditions up until 72h post-transduction and then processed for imaging.

**Flow Microfluidic Assays**

For the flow microfluidics assay, HUVECs were plated at a concentration of $3x10^6$ cells/mL onto iBIDI *μ*-Slide I $^{0.4}$ Luer (iBIDI, 80176), previously coated with 0.2% gelatin solution in $H_2O$ (Sigma-Aldrich, G1393), 4h prior flow application. When transfected cells were used, they were plated 68h after transfection. Flow culture medium consisted of Leibovitz L15 media (Life technologies, LTI 21083-027) supplemented with EGM-2 SingleQuots™ (Lonza, CC-4176) and 1% penicillin/streptomycin (Gibco, #15140122). After 4h of attachment, HUVECs were subjected to flow at different ranges, depending on the experiment, for 4 hours or the appropriate amount of time. The iBIDI slides were connected to a peristaltic pump (Gilson Minipuls3) that ensures a continuous laminar flow during the experiments.

For the flow and scratch wound-assay set of experiments, HUVECs were plated onto microscopy glass slides (Thermo Scientific, 76x26mm) coated with 0.2% gelatin solution in $H_2O$ (Sigma-Aldrich, G1393) at $3x10^6$ cells/mL. When HUVECs reached confluence, a wound was created by scratching the surface of the microscopy glass slide with a 200$\mu$L pipette tip. The culture medium was then replaced by fresh complete medium and HUVECs were allowed to migrate under flow, using sticky-Slide I $^{0.4}$ Luer (iBIDI, 80178) for 5 hours.

**Drugs Assays**

For the experiments with inhibitors, HUVECs were seeded at $3x10^6$ cells/mL in iBIDI $\mu$-Slide I $^{0.4}$ Luer (iBIDI, 80176), previously coated with 0.2% gelatin solution in $H_2O$ (Sigma-Aldrich, G1393), 4h prior flow application. Inhibitors were added to the media 1h before the flow microfluidic assay at specific concentrations. When indicated, cells were treated with (–)-Blebbistatin (20$\mu$M, Sigma-Aldrich, B0560), Y-27632 (5$\mu$M, Merck Millipore, 688001), RGDS (20$\mu$M, Tocris, 3498), Puromycin (200µg/ml, Sigma-Aldrich, P8333) and Triptolide (2µM, Sigma-Aldrich, T3652).

**Immunofluorescence of Cultured HUVECs**

For immunofluorescence of *in vitro* cultured HUVECs, cells were fixed in 1% PFA *Sigma-Aldrich, 4412244) in PBS for 30min at RT. Fixed HUVECs were blocked and permeabilized with blocking solution containing 3% BSA, 0.1% Triton X100 in PBS for 30min at RT. Then cells were incubated for 2h at RT with the appropriate primary antibodies diluted in the blocking solution (anti-VE-cadherin, R&D - AF938, 1:50; anti-GOLPH4, Abcam - ab28049, 1:400; anti-Vinculin, Sigma-Aldrich - V9264, 1:400; anti-ITGA5, Abcam - ab150361, 1:100; anti-pPaxillin, Cell Signaling - 2541S, 1:100; anti-activated ITGB1 BD Pharmingen - 553715, 1:100) and washed 3 x 15min with PBST. Afterwards, cells were incubated in blocking solution containing the secondary fluorophore conjugated antibodies for 1h at RT in the dark (Donkey anti-goat Alexa 647, Thermo Fisher Scientific - A21447, 1:400; Donkey anti-mouse Alexa 488, Thermo Fisher Scientific - A21202, 1:400; Donkey anti-rabbit Alexa 568, Thermo Fisher Scientific - A10042, 1:400), followed again by 3 washes of 15min in PBST. Finally, HUVECs were incubated with 1x DAPI (Molecular Probes by Life Technologies, D1306) diluted in PBS for 5min in the dark. For morphological or colocalization analysis, high-resolution Z-stack images at multiple positions were acquired on a confocal Laser Point-Scanning Microscope 880 (Zeiss) equipped with the Zen black software with a Plan Apochromat 63x NA 1.40 oil DIC M27 objective. For polarization analysis, images at multiple positions were acquired on Zeiss Axiovert 200M equipped with an EC Plan-NeoFluar 40x NA 0.75.

**Protein Extraction and Western Blotting**

Protein extraction was performed from HUVECs lysed in 120$\mu$L of RIPA buffer supplemented with phosphatase and proteinase inhibitors cocktail (1:100, Fischer Scientific, #10085973). Adherent cells were then detached from the plate with a cell scrapper and the cell lysates were transferred into an ice cold eppendorf tube and centrifuged at maximum speed for 10min at 4ºC. Protein concentration was quantified using the BCA protein assay kit (Pierce™, Thermo Fisher Scientific, 23227) following the manufacturer's guidelines. The Multimode microplate reader, Infinite M200 (TECAN), was used for spectrophotometric measurement of protein with the i-control™ software. For Western Blotting, protein samples were normalized up to 25$\mu$L and combined with a mixture of 4x Laemmli Sample Buffer (Bio-rad Laboratories, #161-0747) with 450mM DTT

(SigmaAldrich, D0632) and incubated at 95ºC for 5min. Protein samples were loaded and separated on a 4-15% Mini-PROTEANTGX Gel (BioRad, #456-1084) along with 5μL of protein ladder (GE Healthcare, RPN800E). After transfer, blotted membranes were incubated in Ponceau Red to assess transfer quality, and then washed in TBST (50mM Tris/HCl, 150mM NaCl, 0.1% Tween-20, pH7.5). Afterwards, membranes were incubated in blocking solution containing 3% BSA in TBST for 1h at RT, followed by an o/n incubation at 4°C with the primary antibodies diluted in the same blocking buffer, anti-pPaxillin Y118 (Cell Signalling - 2541S, 1:1000), anti-ßactin (Sigma, A5441, 1:5000), anti-Vinculin (Sigma-Aldrich, V9264, 1:1000), anti-pVinculin Y100 (Thermo Fisher Scientific, 44-1074G, 1:1000), anti-VE-cadherin (Santa Cruz Biotechnology, sc-9989, 1:1000), anti-αCatenin (Sigma-Aldrich, C2081, 1:1000), anti-Talin(Cell Signaling, 4021, 1:1000), anti-AKT (Cell Signaling, 9272, 1:1000), anti-pAKT S473 (Cell Signaling, 4060, 1:1000), anti-FAK (Cell Signaling, 71433, 1:1000), anti-pFAK Y397 (Cell Signaling, 8556, 1:1000), anti-NF-kB p65 (Cell Signaling, 6956, 1:1000) and anti-pNF-kB p65 S536 (Cell Signaling, 3033, 1:1000). On the following day membranes were washed 3 times in TBST and incubated with secondary horseradish peroxidase (HRP)-conjugated antibodies for 1h at RT. Before revelation, membranes were washed again 3 times in TBST for 5min and then incubated in ECL™ Western Blotting Detection Reagent 24 (GE Healthcare, RPN2209) following the manufacturer's protocol. Protein bands were visualized in Amersham 680 (Cytiva) and relative protein quantities were measured using Fiji software[8].

**RNA Extraction and cDNA Synthesis**

RNA extraction was performed from HUVECs using the RNeasy Micro Kit (Qiagen) and the GeneJet RNA Purification Kit (Thermo Scientific) as described by the manufacturer's protocol. RNA concentration was quantified using NanoDrop 1000 (Thermo Scientific) and adjusted equally, followed by DNase I digestion (Thermo Scientific) and cDNA synthesis (Superscript IV First-Strand Synthesis System, Invitrogen). In some cases, a spike-in RNA control was added to the mixture in order to control for gene expression during the real-time quantitative PCR (RT-qPCR). cDNA samples were then diluted in RNAse/DNAse-free water for the subsequent RT-qPCR reactions.

**RT-qPCR**

RT-qPCR was performed using a 7500 Fast Real-Time PCR System (Applied Biosystems) with Power SYBR Green PCR Master Mix (Applied Biosystems) following the standard program of the system previously mentioned. For each reaction, 5μL of cDNA was combined with 10μL of Power SYBR Green PCR Master Mix, 4.5μL of RNAse/DNAse free water and 0.5μL of 4μM primers pool (Forward+Reverse) in a MicroAmp Fast Optical 96-well Reaction Plate (Applied Biosystems).

Primer sequence:

| | | |
|---|---|---|
| GAPDH | Fwd-GTCAAGGCTGAGAACGGGAA; | Rev-TGGACTCCACGACGTACTCA; |
| ACTB | Fwd-CTTCCAGCCTTCCTTCCTGG; | Rev-CAGGGCAGTGATCTCCTTCT; |
| KLF2 | Fwd-GCACGCACACAGGTGAGAAG; | Rev-CCGTGTGCTTTCGGTAGTGG |
| KLF4 | Fwd-GTTCCCATCTCAAGGCACACC; | Rev-GAGCGGGCGAATTTCCATCC; |
| KDR | Fwd-GAACCTCACTATCCGCAGAGT; | Rev-CCAAGTTCGTCTTTTCCTGGGC; |

The expression levels of each sample duplicate were then normalized to GAPDH or spike-in RNA and the $2^{-\Delta\Delta T}$ method was used to calculate relative alterations in gene expression.

**PDMS Gels**

PDMS (Polydimethylsiloxane) was produced with different stiffness. Briefly, Silicone Elastomer (Sylgard 184 Silicone Elastomer, Dow 101697) was mixed with curing agent

(Sylgard 184 Silicone Elastomer, Dow 101697) at three different ratios (by weight): 5:1; 10:1 and 20:1, corresponding to a Young´s elastic modulus of 1000kPa; 580kPa and 280kPa, respectively, followed by degassing in a vacuum chamber for 30min at RT and cured by heating in an oven at 75ºC for 2h, as previously reported[9]. After polymerization, PDMS gels were demolded and sterilized. Afterward, gels were coated with 0.2% gelatin solution in $H_2O$ for 30min at 37ºC and HUVECs seeded as described before for flow experiments.

Soft-PDMS gels with different stiffnesses were produced to measure the focal adhesion length in ECs under shear stress. Briefly, solution A and B (DOWSIL™ CY 52-276 A&B, Dow) were mixed in a 1:1 or 5:6 (w/w) ratio corresponding to a bulk elastic modulus of 3kPa and 18.6 kPa respectively, followed by degassing in a vacuum chamber for 30min on ice. Afterwards, 2ml of soft-PDMS were added to glass slides and spun for 90s, 24V in a handmade spin-coater. Finally, slides with soft-PDMS were cured by heating in an oven at 65ºC o/n.

To promote better cell adhesion, soft-PDMS gels were incubated with 0.2mg/ml of Sulfo-SANPAH (sulfosuccinimidyl 6-(4'-azido-2'-nitrophenylamino) hexanoate, Thermo Scientific) twice during 5 min under UV lamp (approximately 365nm) and washed twice with 50mM of HEPES (Gibco). Finally, soft-PDMS gels were coated with 0.2% gelatin solution in $H_2O$ for 30min at 37ºC and HUVECs seeded 4h prior flow experiments as described previously.

**Traction Force Microscopy and Monolayer Stress Microscopy**

For the TFM experiments, soft-PDMS gels (1:1 ratio) were produced as described previously[9]. To coat with fluorescent beads, 5% APTES (SIGMA, A3648) was used to silanise the soft-PDMS and the gels were washed three times with EtOH absolute. The glass slides with the soft-PDMS were then dried in an oven at 60ºC for 10min. FluoSpheresCarboxylate-Modified Microspheres beads (Invitrogen, F8810) were diluted (1:50) in a boric solution ($Na_2BO_4O_7$, SIGMA 221732 and $H_3BO_3$, SIGMA B1934), filtered with a 0.45µm filter and sonicated for 10min. Afterwards, fluorescent beads were added to the soft-PDMS gels and incubated for 5min at RT. Gels were then dried in an oven at 60ºC. Then, gels were incubated with 0.2mg/ml of Sulfo-SANPAH [sulfosuccinimidyl 6-(4'-azido-2'-nitrophenylamino) hexanoate, Thermo Scientific] twice during 5 min under UV lamp (approximately 365nm) and washed twice with 50mM of HEPES (Gibco). Finally, soft-PDMS gels were coated with 0.2% gelatin solution in $H_2O$ for 30min at 37ºC and HUVECs seeded 4h prior flow experiments.

Traction force measurements were performed as described previously[10]. For each condition, fluorescent images of cell monolayers and nanobeads placed on the surface of the gels were imaged in Leica SP8 multi-photon microscope using the Insight DS+ Dual pulsed laser at 920nm and using a Leica 20x objective (NA 0.95) using tile scans of 4x4 fields-of-view. At the end of the measurements, cells were detached from the gel with 10x trypsin/EDTA (Gibco) and an image of bead position in the relaxed state of the gel was acquired. Individual tiles of each field of view were stitched by using FIJI [8] and the Grid/Collection stitching plugin[11]. 2D images of the deformed substrate were compared to the relaxed state with a custom-made PIV software in Matlab (MathWorks) to obtain the 2D deformation of the top layer of the gel. Fineite-thickness Fourier-transform traction force microscopy was then used to calculate the traction forces exerted by the cells[12,13]. The average forces per unit area exerted by each monolayer were then calculated. To calculate the minimum detectable force levels, we followed the same procedure in cell-free gel areas, and calculated the resulting forces.

Monolayer Stress Microscopy[14,15] was used to calculate the monolayer tension from the traction fields. It was implemented, as a custom-made software, in Python 3 using

NumPy[16], SciPy[17], Matplotlib[18], scikit-image[19], pandas[20], pyFFTW[21], opencv[22] and cython[23].

**Proximity Ligation Assay**

After flow microfluidic experiments, HUVECs were processed for PLA using the Duolink In Situ Red Mouse/Rabbit Starter Kit (Sigma-Aldrich, DUO92101-1KT) as described by the manufacturer's protocol. To probe interactions between VINCULIN and VE-cadherin, cells were incubated with an anti-vinculin antibody raised in rabbit (Sigma-Aldrich, V4139) and an anti-VE-cadherin antibody raised in mouse (Santa Cruz Biotechnologies, sc-9989). In parallel, cells were also incubated with an anti-VE-cadherin antibody raised in goat (R&D Systems, AF938) and subsequently with an anti-goat Alexa 647 secondary antibody (Thermo Fisher Scientific, A21447) to label adherens junctions. To probe interactions between VINCULIN and ITGA5, cells were incubated with an anti-vinculin antibody raised in mouse (Sigma-Aldrich, V9264) and an anti-ITGA5 antibody raised in rabbit (Abcam, ab150361).

To quantify colocalization of PLA signal at adherens junctions, high-resolution Z-stack images at multiple positions were acquired on a confocal Laser Point-Scanning Microscope 880 (Zeiss) equipped with the Zen black software with a Plan Apochromat 63x NA 1.40 oil DIC M27 objective. Briefly, PLA dots were quantified using ImageJ' particle analysis tool and the data normalized by the number of cells.

**Tension Sensor FRET Measurements**

Cells infected with the viral plasmid pRRL-VinculinTS[7] were used for these experiments. FRET images were obtained using a confocal Laser Point-Scanning Microscope 880 (Zeiss) equipped with a Plan-Apochromat 63x, NA 1.40, oil immersion, DIC M27 objective and an argon laser featuring 405, 458 and 514nm laser lines. For FRET experiments, an MBS 458/514 beam splitter was used in combination with the following filters: mTFP1 GaAsP, band-pass 461–520; Venus/FRET, band-pass 525–575. Acceptor photobleaching experiments were analyzed using a custom written MATLAB script. A Gaussian filter with standard deviation of 0.75 was applied to the images before analysis. The intensity in the region of interest was measured before and after bleaching. FRET efficiency was calculated as $EF = \frac{Ipost-Ipre}{Ipost}$, where Ipost and Ipre are the intensity of the donor channel after and before bleaching respectively.

**Colocalization Analysis**

For colocalization analysis, high-resolution Z-stack confocal images of HUVECs stained for different proteins (VE-Cadherin, Vinculin and ITGA5) were imported and analyzed in MATLAB using a custom written code. An object-based co-localization approach was performed. Concisely, each channel was segmented, and a binary mask generated. The masks were combined and the fraction of pixels with overlapping signals was quantified.

**Analysis of adhesion orientation and vinculin-aITGB1 colocalization angle**

To analyze FA orientation, high resolution Z-stack confocal images of HUVECs stained for vinculin and aITGB1 were segmented and binarized using ImageJ and then imported and analyzed in MATLAB using a custom written code. An object-based analysis approach was performed. Briefly, each vinculin object orientation was computed to determine FA adhesion orientation. Each vinculin and aITGB1 object centroid coordinated were determined. Each vinculin object was paired to the closest aITGB1 object, and their respective centroids used to determine a vinculin-to-aITGB1 vector. Vector angles were computed to determine the polarization of aITGB1 related to vinculin.

**Vascular Morphometrics Analysis**

For radial expansion quantification, images were taken using an EC Plan-Neofluar, NA 0.30, air 10x objective, in a confocal Laser Point-Scanning Microscope 880 (Zeiss). The migratory length of the vascular plexus was analyzed by measuring the total length of the retinal vasculature from the optic nerve, in the center, towards the retinal periphery – sprouting front.

For the quantification of the number of tip cells, tile-scan images of the whole sprouting front were taken with a C-Apochromat Corr, NA 1.20, water 40x objective in a confocal Laser Point-Scanning Microscope 880 (Zeiss). The number of tip cells was then counted, and the values normalized by the sprouting front length.

Regarding vessel density, tile-scan images of the whole petal were acquired using a Plan-Apochromat, NA 0.8, air 20x objective in a confocal Laser Point-Scanning Microscope 880 (Zeiss). After vessel segmentation using FIJI and Photoshop, the area of the vessels was calculated using a MATLAB script developed in our lab. Vessel density was calculated as a ratio between vessel area and total area of the petal.

Endothelial cell density was calculated using 20x tile-scan images acquired in a confocal Laser Point-Scanning Microscope 880 (Zeiss). The number of $ERG^+$ nuclei counted manually was normalized by the vascularized area obtained from the MATLAB script for each petal (two petals for each retina).

**Vascular Morphogenesis Parameters and Principal Component Analysis**

Retinal vascular plexuses were imaged, binarized, and skeletonized following the protocol previously described in Bernabeu et al. 2018[24]. Vessel diameter was stored as a node attribute in the graph data structure. The resulting planar graphs were manually cropped to the arterial region of interest and simplified by merging edges that met at nodes of degree 2. Every remaining vertex in the graph was therefore a bifurcation (degree 3), the tip of blind-ended vessel (sprout or vessel undergoing pruning) (degree 1) or vessel leaving the region of interest (degree 1). The faces of the planar graph were obtained, and their area calculated. The sprouting front boundary was defined as the line connecting the tips of the two sprouts protruding the most in the sprouting front. With this line as reference, the following graph properties were calculated in bins of 100 $\mu$m width moving away from the sprouting front: vessel density (number of graph edges per unit of vascularized area in the bin), bifurcation density (number of degree 3 nodes per unit of vascularized area in the bin), range of avascular areas (difference between maximum and minimum area of the faces calculated in the bin), mean avascular area (mean of the distribution of face areas in the bin), standard deviation of avascular areas (standard deviation of the distribution of face areas in the bin), mean diameter (mean of the distribution of vessel diameters in the bin), and standard deviation of diameters (standard deviation of the distribution of vessel diameters in the bin). Principal component analysis (PCA) decomposition of the 7-dimensional vectors defining the previous features at each bin in each control retina was performed to facilitate visualization and bin classification based on phenotypic similarity. Weights were as following: PCA1 [-0.453; -0.345; 0.348; 0.366; 0.404; -0.406; -0.304]; PCA2 [0.346; 0.0638; 0.523; 0.220; 0.491; 0.423; 0.366], for vessel density; bifurcation density; range of avascular areas; mean avascular area; standard deviation of avascular areas; mean diameter; and standard deviation of diameters, respectively. The k-means clustering algorithm was used to find two clusters that minimize within cluster variance under the assumption that two main phenotypic classes (sprouting and remodeling) exist. The PCA decomposition of the control group was used to map the bins of the remaining groups to the same phenotypic space. The k-means classifier was used to classify these bins according to their distance to the center of the sprouting/remodeling clusters in the control group.

**Polarity Index Quantifications (*in vitro*)**
To quantify cell polarity, tile-scan images of HUVECs stained for Golgi (Golph4) and nuclei (DAPI) markers were processed in FIJI. Afterwards, each set of images was imported and analyzed in MATLAB using a modified version of a polarity analysis script kindly provided by Anne-Clémence Vion and Holger Gerhardt.

Briefly, after segmenting each channel corresponding to the Golgi and nuclear staining, the centroid of each organelle was determined and a vector connecting the center of the nucleus to the center of its corresponding Golgi apparatus was drawn. The Golgi-nucleus assignment was done automatically minimizing the distance between all the possible couples. The polarity of each cell was defined as the angle between the vector and the slide axis. An angular histogram showing the angle distribution was then generated. Circular statistics were performed using the Circular Statistic Toolbox. The polarity index (PI) was calculated as the length of mean resultant vector for a given angular distribution.

$$\text{Polarity Index} = \sqrt{\left(\frac{1}{N}\sum_{1}^{N}\cos\alpha\right)^2 + \left(\frac{1}{N}\sum_{1}^{N}\sin\alpha\right)^2}$$

The PI varies between 0 and 1. The closer to 1, the more the data are concentrated around the mean direction, while values close to 0 correspond to random distribution. PI indicates the collective orientation strength of the cell monolayer. Box plots were generated by using every single PI calculated for images of each biological replica, which show the biological variability of the system. This data was used to calculate the significance of differences between experimental conditions.

**Flow and Chemokine Polarity Indexes in Retinas**
In the *in vivo* polarity analysis, we quantified polarity and correlated it with blood flow direction by using the approach described in Bernabeu et al. 2018[24]. Briefly, all the retinal vascular plexuses were imaged in a Zeiss Cell Observer SD (Zeiss) equipped with Zen software and with a Plan-Apochromat 40x NA 1.40 oil objective and post-processed to generate a binary mask from the ICAM2 channel and a second image with at least the ERG (EC nuclei) and Golph4 (Golgi apparatus) channels. These two images defined the input to PolNet. PolNet is a graphical user interface that allows the user to perform three tasks: 1) to construct a flow model from the ICAM2 mask and use the HemeLB flow solver to estimate the wall shear stress across the whole network (as well as blood velocity, shear rate, and pressure); 2) to draw cell polarity vectors (nuclei to Golgi) for each endothelial cell in the network based on the Erg-Golph4 image; and 3) to statistically analyze the relationship between the cell polarity and flow direction or wall shear stress.

Chemokine (K-) and Flow (F-) polarity indexes were then calculated using the angle that each Nuclei-to-Golgi vector does with either the sprouting front edge (K-), defined as (defined as a line between the two most outward vascular sprouts), or the flow direction (F-) using the length of mean resultant vector for a given angular distribution, as previously described[24-26]. The polarity index varies between -1 (backward polarization – away from the sprouting front, if K-index; or with the flow direction, if F-index) to 1 (forward polarization – towards free edge of the sprouting front, if K-index; or against the flow direction, if F-index), where 0 means random polarization. Chemokine (K-) and Flow (F-) polarity indexes were then represented as a function of the distance from the sprouting front (SF) to the optic nerve (ON).

**Statistical Analysis**
All statistical analysis was performed using GraphPad Prism 7. Measurements were taken from distinct samples, and statistical details of experiments are reported in the figures and figure legends. Sample size is reported in the figure legends and no statistical test was used to determine sample size. The biological replicate is defined as the number of cells, images, animals, as stated in the figure legends. No inclusion/exclusion or randomization criteria were used and all analyzed samples are included. Normality tests were performed to assess the data normality. Comparisons between two experimental groups were analyzed with two-sided unpaired parametric t test or Mann-Whitney test depending on the data normality. Multiple comparisons between more than two experimental groups were assessed with one-way ANOVA. We considered a result significant when $p<0.05$. For all box plots: centerline, median; +, mean; whiskers, min to max.

#### Supplementary References


1 Leon, C. *et al.* Megakaryocyte-restricted MYH9 inactivation dramatically affects hemostasis while preserving platelet aggregation and secretion. *Blood* **110**, 3183-3191, doi:10.1182/blood-2007-03-080184 (2007).
2 Vasioukhin, V., Bauer, C., Degenstein, L., Wise, B. & Fuchs, E. Hyperproliferation and defects in epithelial polarity upon conditional ablation of alpha-catenin in skin. *Cell* **104**, 605-617, doi:10.1016/s0092-8674(01)00246-x (2001).
3 Claxton, S. *et al.* Efficient, inducible Cre-recombinase activation in vascular endothelium. *Genesis* **46**, 74-80, doi:10.1002/dvg.20367 (2008).
4 Sörensen, I., Adams, R. H. & Gossler, A. DLL1-mediated Notch activation regulates endothelial identity in mouse fetal arteries. *Blood* **113**, 5680-5688, doi:10.1182/blood-2008-08-174508 (2009).
5 Figueiredo, A. M. *et al.* Endothelial cell invasion is controlled by dactylopodia. *Proc Natl Acad Sci U S A* **118**, doi:10.1073/pnas.2023829118 (2021).
6 Franco, C. A. *et al.* SRF selectively controls tip cell invasive behavior in angiogenesis. *Development* **140**, 2321-2333, doi:10.1242/dev.091074 (2013).
7 Rothenberg, K. E., Scott, D. W., Christoforou, N. & Hoffman, B. D. Vinculin Force-Sensitive Dynamics at Focal Adhesions Enable Effective Directed Cell Migration. *Biophys J* **114**, 1680-1694, doi:10.1016/j.bpj.2018.02.019 (2018).
8 Schindelin, J. *et al.* Fiji: an open-source platform for biological-image analysis. *Nature Methods* **9**, 676-682, doi:10.1038/nmeth.2019 (2012).
9 Park, J. Y. *et al.* Increased poly(dimethylsiloxane) stiffness improves viability and morphology of mouse fibroblast cells. *BioChip Journal* **4**, 230-236, doi:10.1007/s13206-010-4311-9 (2010).
10 Butler, J. P., Tolić-Nørrelykke, I. M., Fabry, B. & Fredberg, J. J. Traction fields, moments, and strain energy that cells exert on their surroundings. *American journal of physiology. Cell physiology* **282**, C595-605, doi:10.1152/ajpcell.00270.2001 (2002).
11 Preibisch, S., Saalfeld, S. & Tomancak, P. Globally optimal stitching of tiled 3D microscopic image acquisitions. *Bioinformatics* **25**, 1463-1465, doi:10.1093/bioinformatics/btp184 %J Bioinformatics (2009).
12 Trepat, X. *et al.* Physical forces during collective cell migration. *Nature Physics* **5**, 426-430, doi:10.1038/nphys1269 (2009).
13 Elosegui-Artola, A. *et al.* Rigidity sensing and adaptation through regulation of integrin types. *Nature materials* **13**, 631-637, doi:10.1038/nmat3960 (2014).
14 Tambe, D. T. *et al.* Monolayer Stress Microscopy: Limitations, Artifacts, and Accuracy of Recovered Intercellular Stresses. *PLOS ONE* **8**, e55172, doi:10.1371/journal.pone.0055172 (2013).

15 Tambe, D. T. *et al.* Collective cell guidance by cooperative intercellular forces. *Nature materials* **10**, 469-475, doi:10.1038/nmat3025 (2011).

16 Harris, C. R. *et al.* Array programming with NumPy. *Nature* **585**, 357-362, doi:10.1038/s41586-020-2649-2 (2020).

17 Virtanen, P. *et al.* SciPy 1.0: fundamental algorithms for scientific computing in Python. *Nature Methods* **17**, 261-272, doi:10.1038/s41592-019-0686-2 (2020).

18 Hunter, J. D. Matplotlib: A 2D Graphics Environment. *Computing in Science & Engineering* **9**, 90-95, doi:10.1109/MCSE.2007.55 (2007).

19 van der Walt, S. *et al.* scikit-image: image processing in Python. *PeerJ* **2**, e453, doi:10.7717/peerj.453 (2014).

20 McKinney, W. Data Structures for Statistical Computing in Python. *Proceedings of the 9th Python in Science Conference*, 56 - 61, doi:10.25080/Majora-92bf1922-00a (2010).

21 Frigo, M. in *Proceedings of the ACM SIGPLAN 1999 conference on Programming language design and implementation* 169–180 (Association for Computing Machinery, Atlanta, Georgia, USA, 1999).

22 Bradski, G. The OpenCV Library. *Dr. Dobb's Journal of Software Tools* **25**, 120-125 (2000).

23 Behnel, S. *et al.* Cython: The Best of Both Worlds. *Computing in Science & Engineering* **13**, 31-39, doi:10.1109/MCSE.2010.118 (2011).

24 Bernabeu, M. O. *et al.* PolNet: A Tool to Quantify Network-Level Cell Polarity and Blood Flow in Vascular Remodeling. *Biophys J* **114**, 2052-2058, doi:10.1016/j.bpj.2018.03.032 (2018).

25 Carvalho, J. R. *et al.* Non-canonical Wnt signaling regulates junctional mechanocoupling during angiogenic collective cell migration. *Elife* **8**, doi:10.7554/eLife.45853 (2019).

26 Franco, C. A. *et al.* Dynamic endothelial cell rearrangements drive developmental vessel regression. *PLoS Biol* **13**, e1002125, doi:10.1371/journal.pbio.1002125 (2015).

#### Figure S1

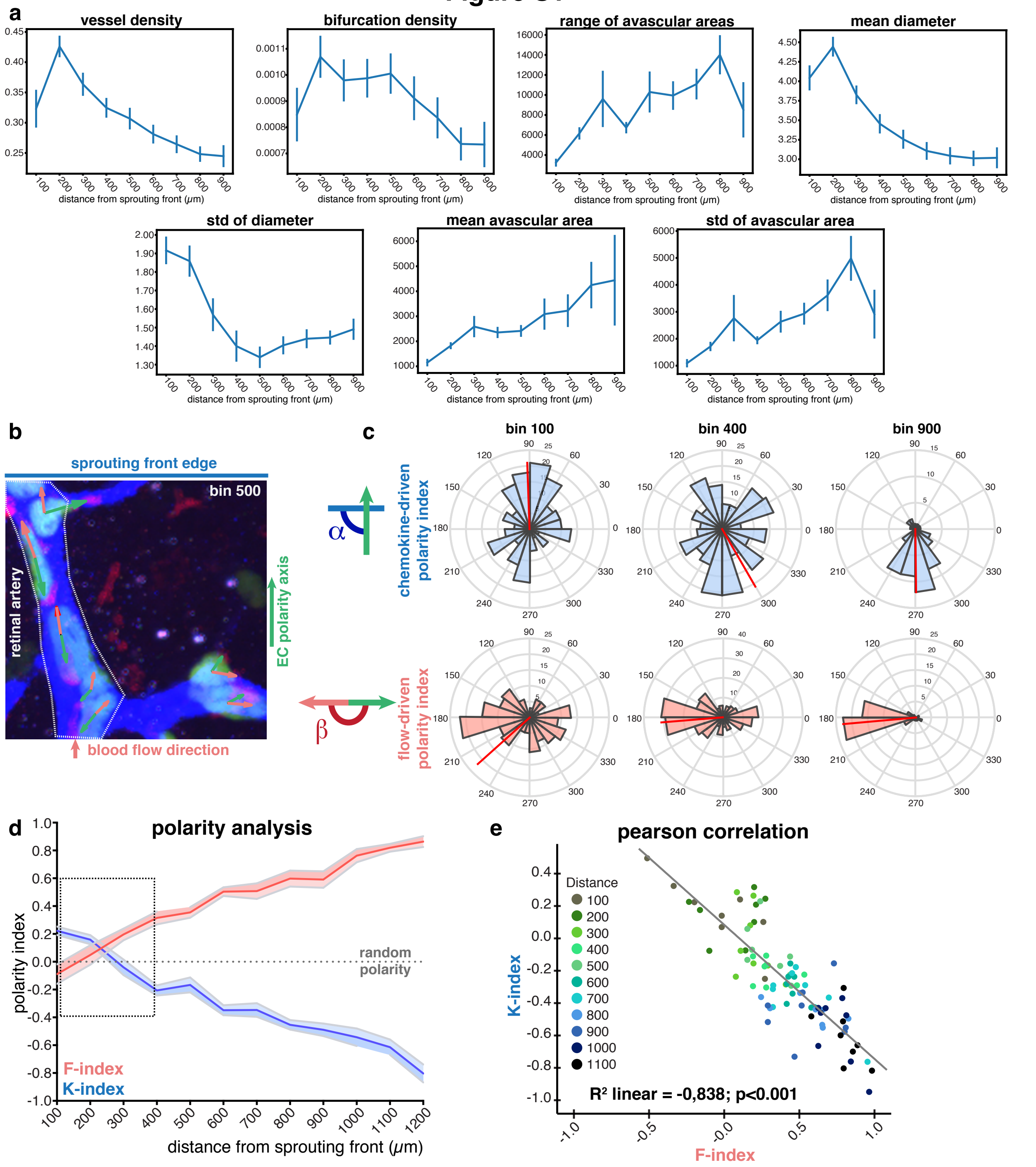


**Figure S1. Vessel morphometric parameters and polarization patterns.**
**a**, Distribution of each vessel morphometric parameter along the sprouting front-to-optic nerve (SF>ON) axis of segmentation for control retinas. **b**, Representative image of the EC polarity (green), sprouting front edge (blue) and blood flow direction (red) used for the calculation of chemokine-induced (K) and flow-induced (F) angles. EC nucleus (ERG, green); vessel lumen (ICAM2, blue); Golgi complex (GOLPH4, red). **c**, Representation of angular histograms for bin 100, bin 400 and bin 900 of K-index (light blue) and F-index (light red) in the vascular network along the SF>ON axis of segmentation. **d**, Distribution of K- (blue) and F-indexes (red) along the SF>ON axis. Solid line represents mean, and light area represents SEM. Dashed black line represents random polarity. Dashed black rectangle highlights zoomed section showed in fig. 2D. N= 11 arterial regions. **e,** Pearson correlation analysis between K- and F-indexes distributed along the SF>ON axis of segmentation. Each dot represents one bin in each retina, and it has been color-coded for the corresponding bin number.

#### Figure S2

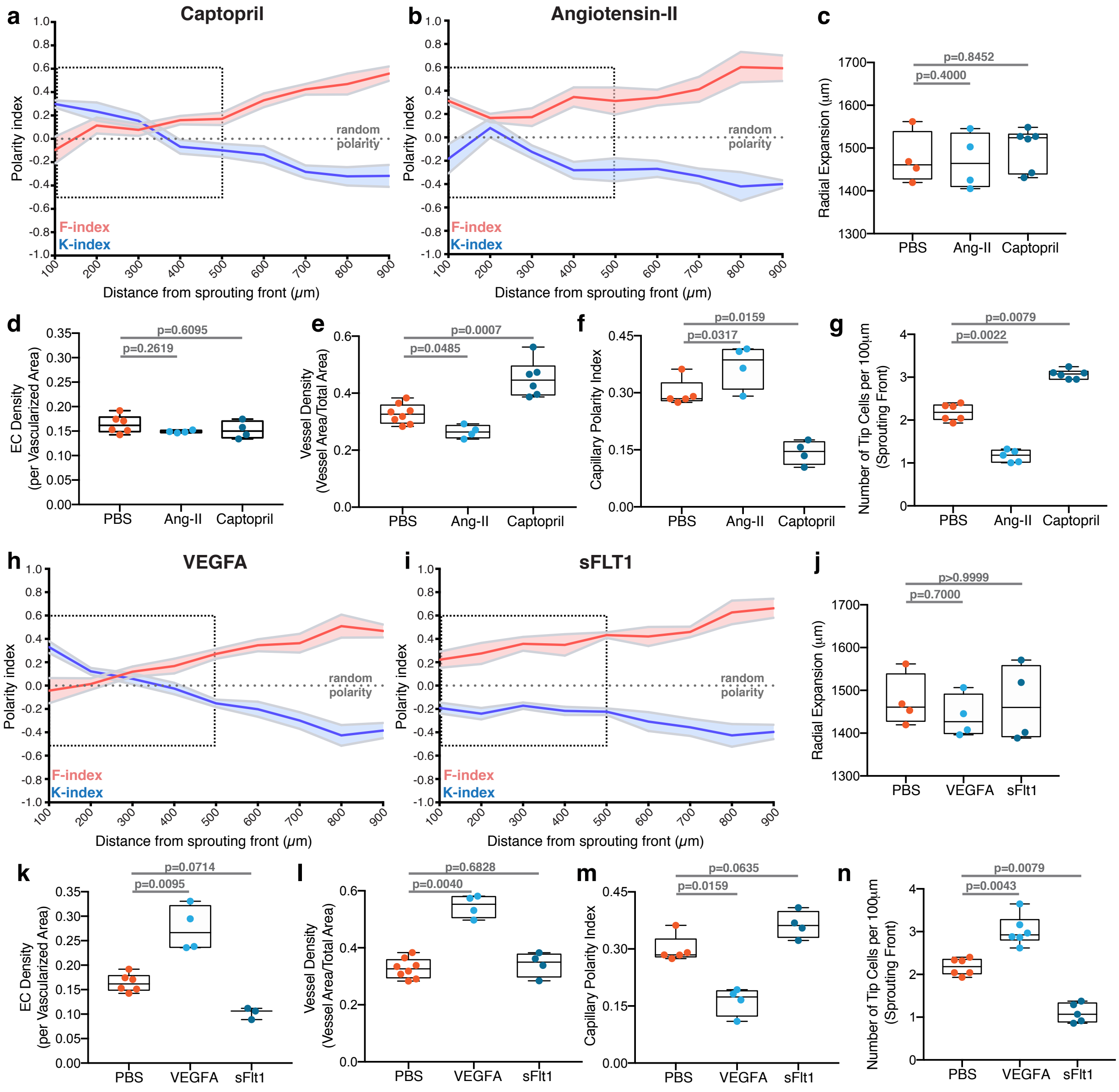


**Figure S2. VEGFA and flow patterns govern the S>R transition.**
**a**, Distribution of K-indexes (blue) and F-indexes (red) along the sprouting front to optic nerve axis of segmentation in Captopril injected mice. Solid line represents mean and light area represents SEM. Dashed black line represents random polarity pattern. Dashed black rectangle highlights zoomed section showed in fig. 3B. N= 4 arterial regions. **b**, Distribution of K-indexes (blue) and F-indexes (red) along the sprouting front to optic nerve axis of segmentation in Angiotensin-II injected mice. Solid line represents mean, and light area represents SEM. Dashed black line represents random polarity pattern. Dashed black rectangle highlights zoomed section showed in fig. 2D. N= 7 arterial regions. **c**, Boxplot for quantification of radial expansion (in μm) for PBS (N= 4), Ang II (N= 4) and Captopril (N= 6) injected pups; p-values from Mann-Whitney test. **d**, Box plot for quantification of EC density per vascularized area for PBS (N= 6), Ang II (N= 4) and Captopril (N= 4) injected pups. p-values from Mann-Whitney test. **e**, Boxplot for quantification of vessel density (vessel area/total area) for PBS (N= 8), Ang II (N= 4) and Captopril (N= 6) injected pups. p-values from Mann-Whitney test. **f**, Box plot for Capillary Polarity Index for PBS (N= 5), Ang II (N= 4) and Captopril (N= 4) injected pups. p-values from Mann-Whitney test. **g**, Boxplot for Number of tip cells per 100µm of the SF (Sprouting Front) for PBS (N= 6), Ang II (N= 5) and Captopril (N= 6) injected pups. p-values from Mann-Whitney test. **h**, Distribution of K-indexes (blue) and F-indexes (red) along the sprouting front to optic nerve axis of segmentation in VEGFA injected mice. Solid line represents mean, and light area represents SEM. Dashed black line represents random polarity

pattern. Dashed black rectangle highlights zoomed section showed in fig. 3F. N= 9 arterial regions. **i**, Distribution of K-indexes (blue) and F-indexes (red) along the sprouting front to optic nerve axis of segmentation in sFLT1 injected mice. Solid line represents mean, and light area represents SEM. Dashed black line represents random polarity pattern. Dashed black rectangle highlights zoomed section showed in fig. 3H. N= 4 arterial regions. **j**, Boxplot for quantification of radial expansion (in $\mu$m) for PBS (N= 4), VEGFA (N= 4) and sFLT1 (N= 4) injected pups. p-values from Mann-Whitney test. **k**, Box plot for quantification of EC density per vascularized area for PBS (N= 6), VEGFA (N= 4) and sFLT1 (N= 3) injected pups. p-values from Mann-Whitney test. **l**, Box plot for quantification of vessel density (vessel area/total area) for PBS (N= 8), VEGFA (N= 4) and sFLT1 (N= 4) injected pups. P-values from Mann-Whitney test. **m**, Box plot for Capillary Polarity Index for PBS (N= 5), VEGFA (N= 4) and sFLT1 (N= 4) injected pups. p-values from Mann-Whitney test. **n**, Box plot for Number of tip cells per 100µm of the SF (Sprouting Front) for PBS (N= 6), VEGFA (N= 6) and sFLT1 (N= 5) injected pups. p-values from Mann-Whitney test.

### Figure S3

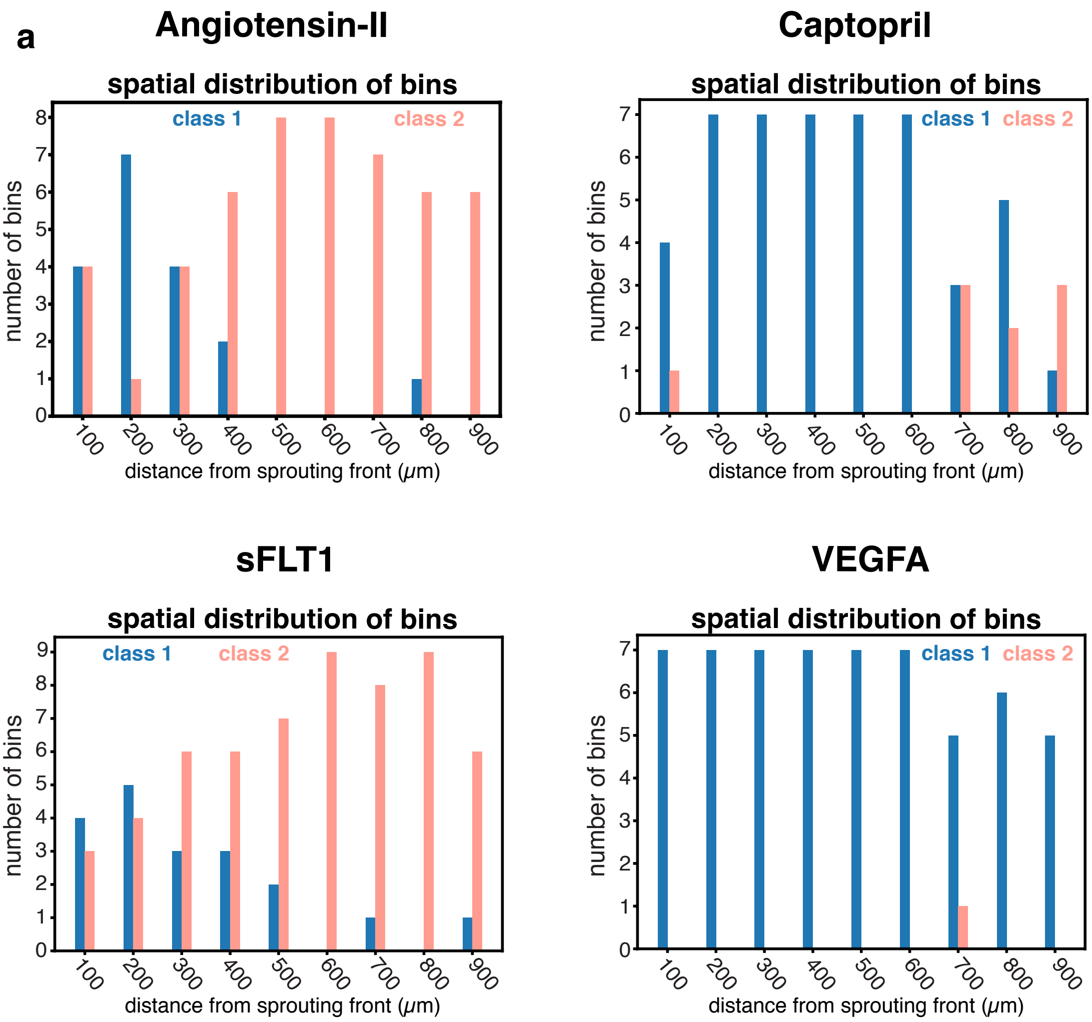


**Figure S3. Disrupted S>R transition in pharmacological perturbations on blood flow and VEGFA levels.**
**a,** Distribution of the number of bins along the SF>ON axis of segmentation and color-coded according to the PCA classification (class 1 or class 2), in angiotensin-II, captopril, sFLT1 or VEGFA treated animals.

### Figure S4

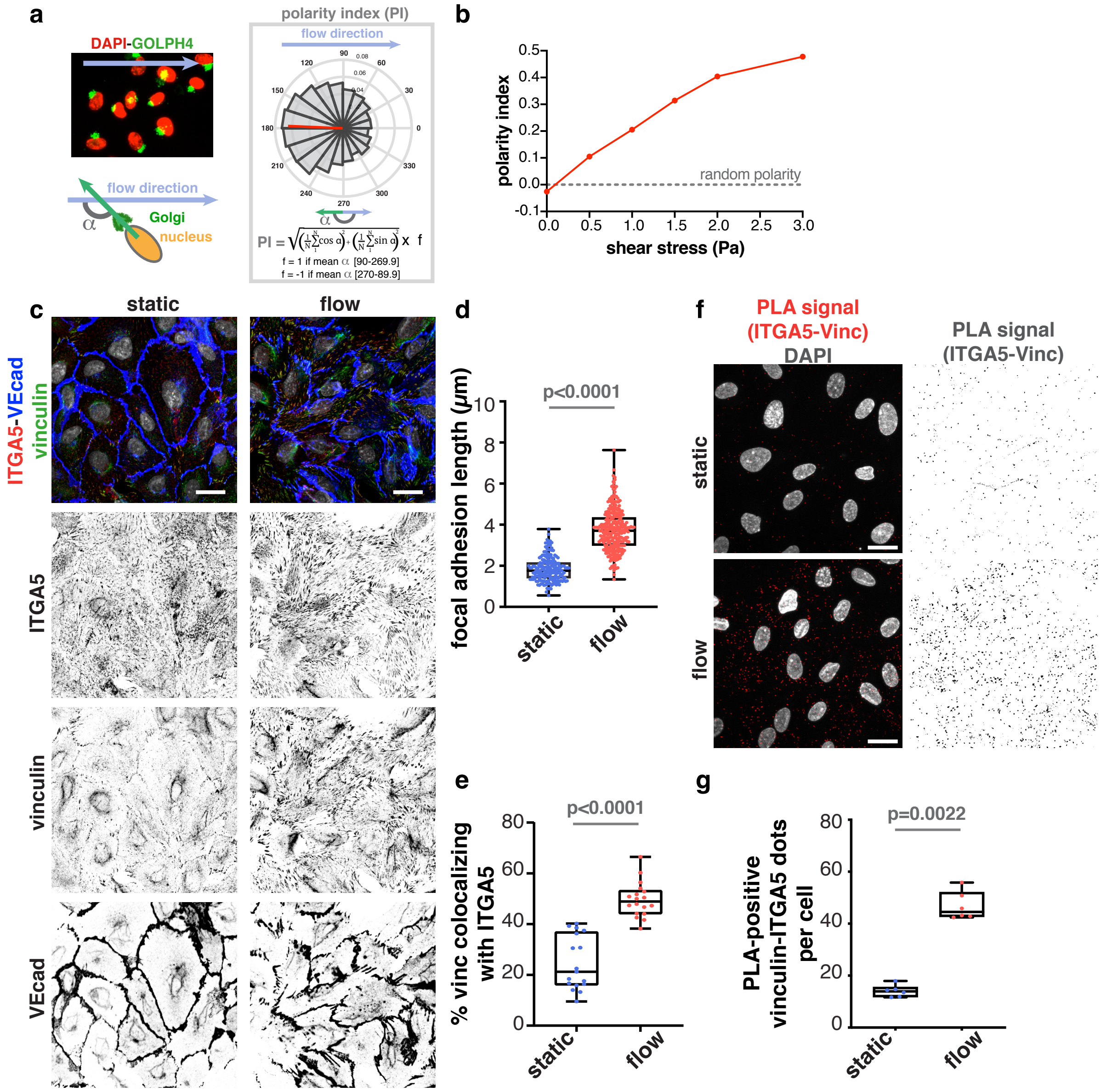


**Figure S4. Flow-induced shear stress strengths focal adhesion-mediated attachment to the extracellular matrix.**

**a**, Top left: representative image of HUVECs polarizing against the flow direction (blue arrow) in the microfluidic chamber. Bottom left: representation of the endothelial cell angle of polarity (α), defined by the angle for nucleus to Golgi apparatus axis in relation to the blood flow direction, flow-induce angle. Right: Calculation of the polarity index (PI) based on α. **b,** Polarity index of HUVEC monolayers exposed to different flow-induced shear stress levels. n= 1 experiment per condition. **c,** Representative image of adherens junctions (VE-cadherin – VEcad, blue) and focal adhesions (vinculin, green, and integrin alpha 5 – ITGA5, red) in static or flow-stimulated HUVEC monolayers (lower-magnification of fig. 4A). Scale bar: 20 µm. Flow direction is right-to-left. **d,** Box plot for quantification of length of focal adhesions in static or flow conditions. For the focal adhesion length, N= 150 per condition; p-values from Mann-Whitney test. **e**, Box plot for quantification of percentage of vinculin colocalizing with integrin alpha 5 (ITGA5) in static or flow conditions. N= 18 per condition (for colocalization); p-values from Mann-Whitney test. **f**, Representative image of HUVEC monolayers under static or flow (2.0 Pa) conditions with nuclei (DAPI, in grey) and Proximity Ligation Assay (PLA) signal (ITGA5-Vinculin, in red). Scale bar: 20µm. **g**, Box plot for quantification of percentage of PLA-positive vinculin-ITGA5 dots in static or flow conditions. N= 6 per condition (PLA); p-values from Mann-Whitney test.

### Figure S5

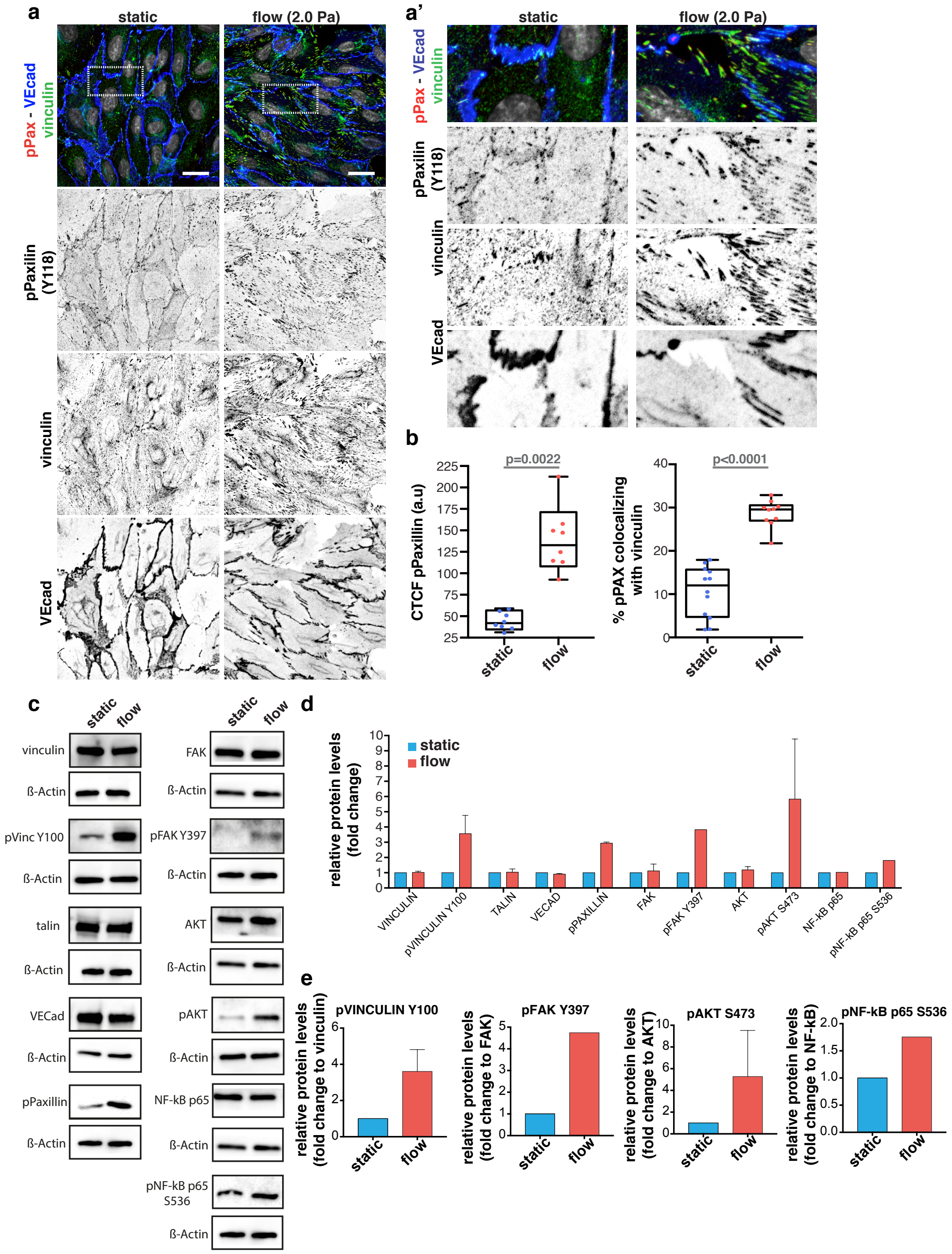


**Figure S5. Flow-induced shear stress activates FA-mediated signaling pathways.**
**a**, Representative image of HUVEC monolayers under static or flow (2.0 Pa) conditions with adherens junction (VEcad, in blue) and focal adhesions (Vinculin, in green and pPaxillin Y118, in red). White square highlights the high magnification zoomed image of A'. Scale bar: 20µm. **b,** Box plot for quantification of corrected total cell fluorescence (CTCF) for phosphorylated paxillin (pPaxillin), and percentage of pPaxillin colocalizing with vinculin in static or flow conditions. N= 8 (for CTCF); N= 11 (for static colocalization) and 10 (for flow colocalization); p-values from Mann-Whitney test. **c**, Western blots for several proteins from HUVECs under static or flow conditions. **d**, Quantification of protein levels normalized to b-actin (static conditions and flow conditions). N= 2. **e,** Quantification of pVinculin Y100 levels normalized to total Vinculin under static or flow conditions; N= 2. Quantification of pFAK Y397 levels normalized to total FAK under static or flow conditions; N= 1. Quantification of pAKT S473 levels normalized to total AKT under static or flow conditions; N= 2. Quantification of pNF-kB p65 S536 levels normalized to total NF-kB p65 under static or flow conditions; N= 1.

#### Figure S6

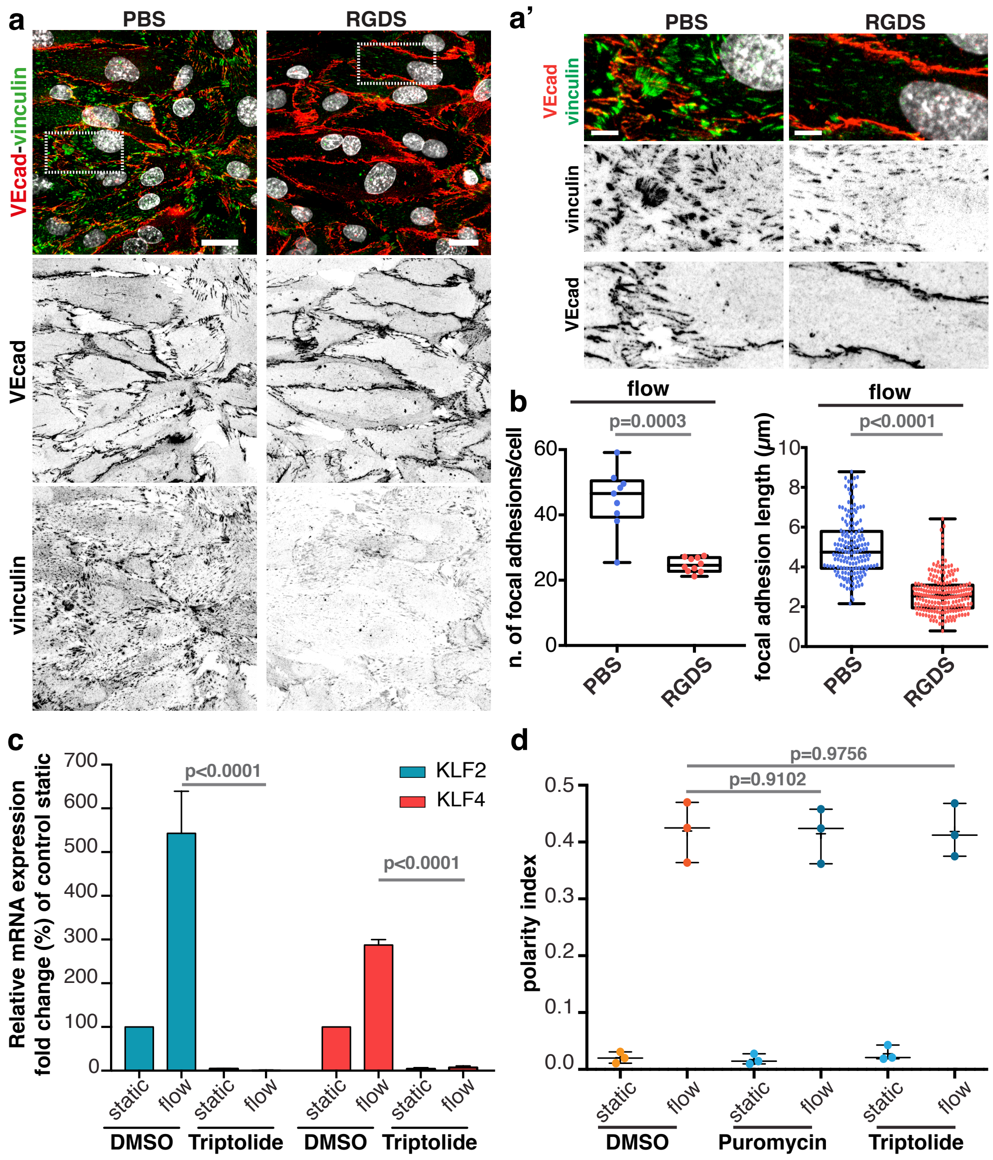


**Figure S6. RGDS treatment reduces focal adhesions and flow-induced polarity is transcription- and translation-independent.**
**a**, Representative image of HUVEC monolayers under flow (2.0 Pa) conditions treated with PBS or RGDS and labelled for adherens junctions (VEcad, in red) and focal adhesions (Vinculin, in green). Scale bar: 20 $\mu$m. White square highlights the high magnification zoomed image of A'. Scale bar: 5µm. **b**, Box plot for quantification of number and length of focal adhesions in flow-stimulated HUVEC monolayers treated with PBS or RGDS. N= 10 per condition (for n of focal adhesions/cell) and N= 200 per condition (for focal adhesion length); p-values from Mann-Whitney test (number of focal adhesions) and from unpaired t test (focal adhesion length). **c**, Quantification of relative mRNA expression (Fold Change % for DMSO static) for flow-responsive genes (Klf2 and Klf4) under static and flow conditions for DMSO and triptolide treated HUVECs. N= 3 per condition. p-values from Mann-Whitney test. **d**, Boxplot of static or flow-induced polarity index in DMSO, puromycin or triptolide HUVEC monolayers. N= 3 per condition. p-values from Mann-Whitney test.

### Figure S7

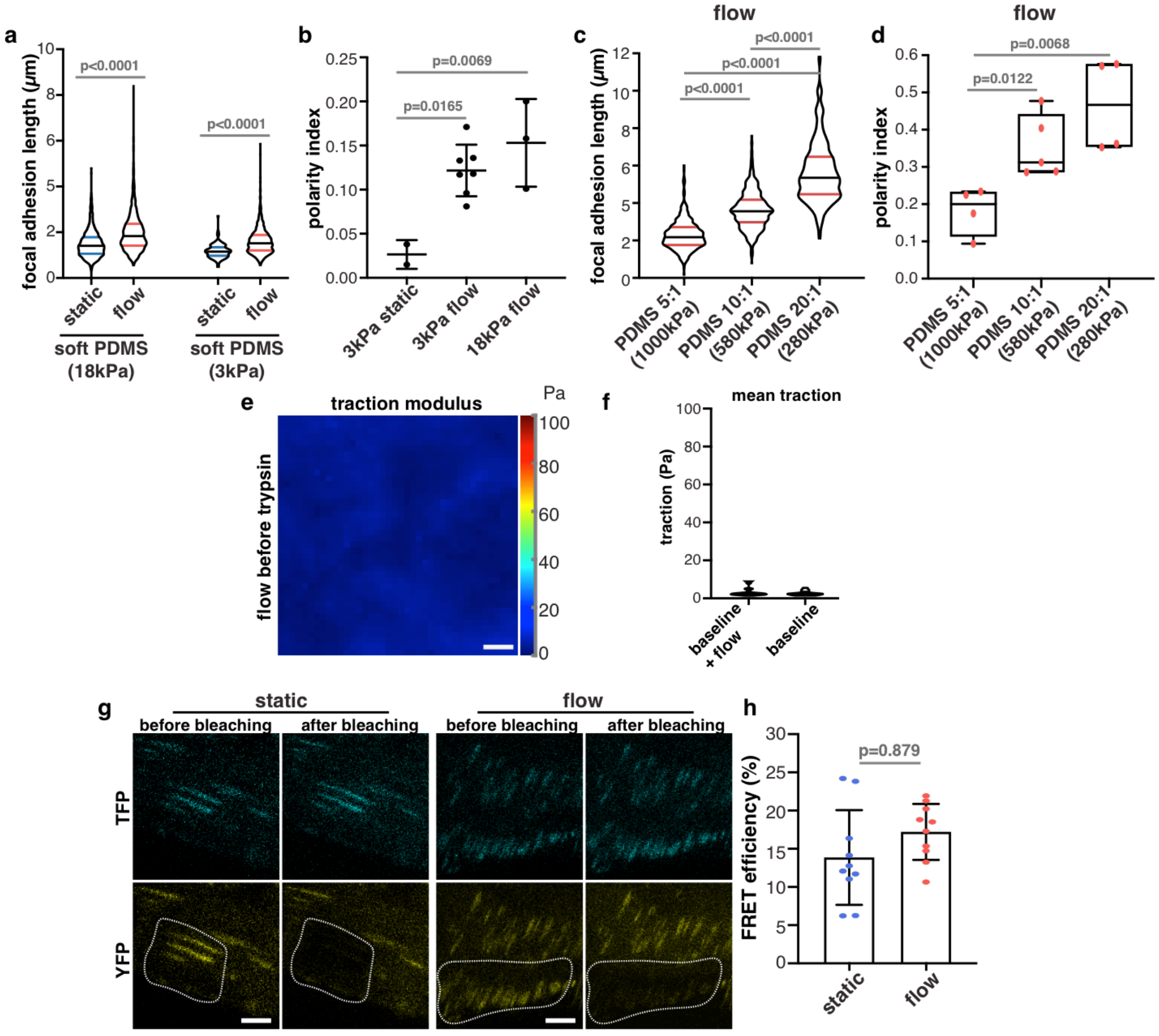


**Figure S7. Shear stress-induced polarity depends on FA-mediated traction forces.**
**a**, Violin plot for quantification of focal adhesion length in flow-stimulated HUVEC monolayers seeded on soft PDMS of 3kPa and 18kPa stiffnesses. N> 200 focal adhesions per condition; p-values from unpaired t test. **b**, Scatter dot plot quantification of polarity index in flow-stimulated HUVEC monolayers seeded on soft PDMS of 3kPa and 18kPa stiffnesses. N= 2 static 3kPa; N= 5 flow 3kPa; N= 3 flow 18kPa; p-values from unpaired t test. **c**, Violin plot for quantification of focal adhesion length in flow-stimulated HUVEC monolayers seeded on PDMS with designated stiffnesses. N> 400 per condition; p-values from unpaired t test. **d**, box plot for polarity index in flow-stimulated HUVEC monolayers seeded on PDMS with designated stiffnesses. N= 4 (5:1 and 20:1); N=5 (10:1); p-values from unpaired t test. **e**, Representative mean traction maps in experimental conditions without cells, following the same protocol as for experiments with cells, to assess the influence of flow in traction forces. **f**, Quantification of mean traction forces in experimental conditions without cells, following the same protocol as for experiments with cells, to assess the influence of flow in traction forces. Quantification shows the effect of taking sequential images before and after trypsin/EDTA addition (which in real experiments is used to remove cells, baseline), and the added effect of flow (by comparing images with and without flow). N= 2 independent experiments. **g**, Representative images of acceptor (YFP, yellow) and donor (TFP, cyan) fluorophores before and after photo-bleaching. White dotted line represents bleached region. Scale bar: 5$\mu$m. **h**, Bar graph for FRET efficiency (in %) of vinculin tension-sensor in static or flow-stimulated HUVEC monolayers. N= 10 per condition; p-values from Mann-Whitney test.

### Figure S8

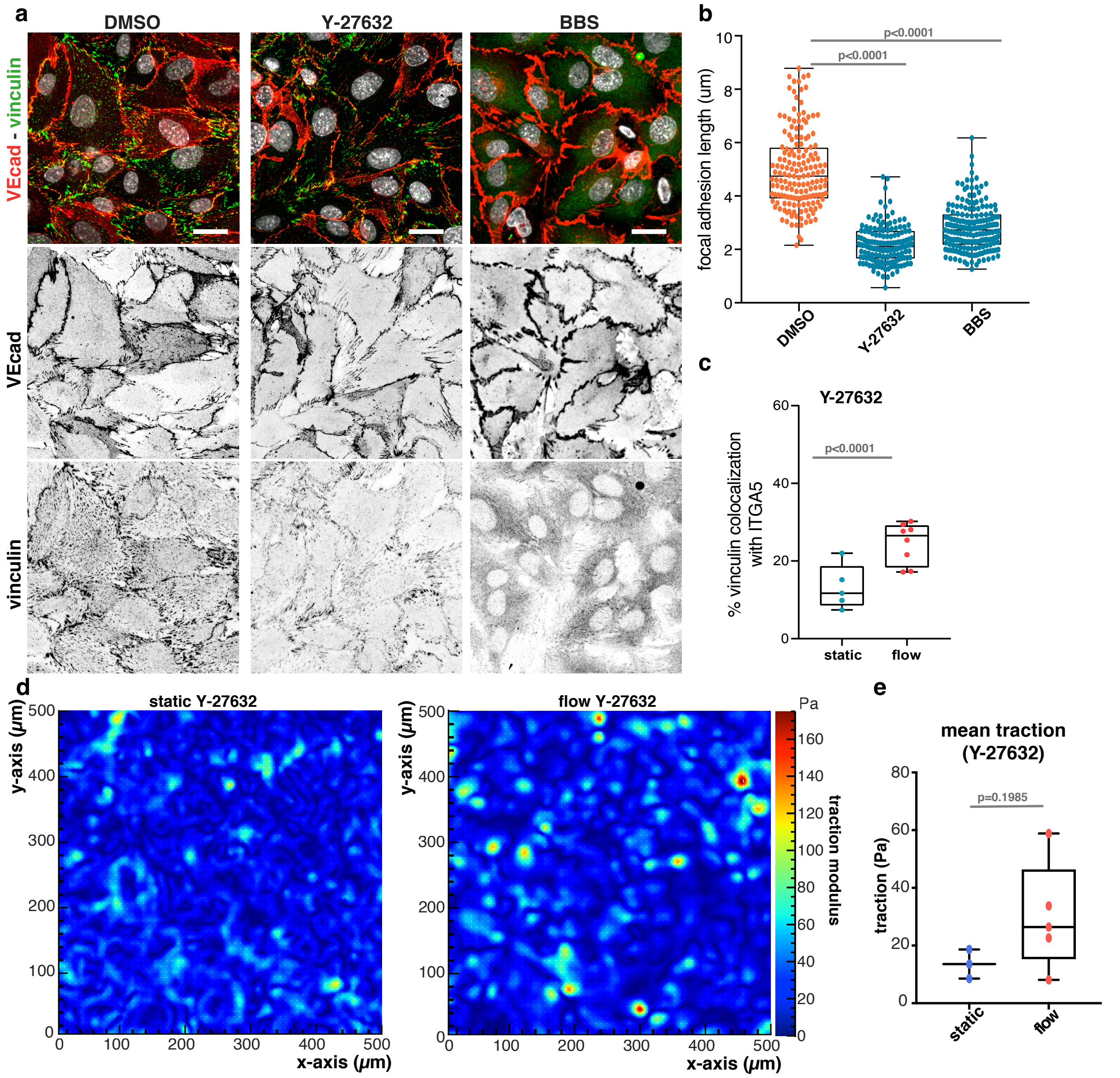


**Figure S8. Inhibition of actomyosin-mediated contractility reduces numbers focal adhesions.**

**a**, Representative image of HUVEC monolayers under flow (2.0 Pa) conditions treated with DMSO, Y-27632 or BBS, labelled for adherens junctions (VEcad, in red) and focal adhesions (Vinculin, in green). Scale bar: 20um. Flow direction is right-to-left. **b**, Boxplot for quantification of focal adhesion length (μm) in flow-stimulated HUVEC monolayers treated with DMSO, Y-27632 or BBS. N= 180 per condition. p-values from unpaired t test. **c**, Boxplot for quantification of % of vinculin colocalization with ITGA5 in HUVEC monolayers treated with Y-27632. N= 5 (static) and N=8 (flow); p-values from Mann-Whitney test. **d**, Representative mean traction maps exerted by static or flow-stimulated HUVEC monolayers treated with Y-27632. **e**, Box plot of mean traction forces exerted by static or flow-stimulated HUVEC monolayers treated with Y-27632. N= 3 static; 5 flow independent experiments; p-values from Mann-Whitney test.

### Figure S9

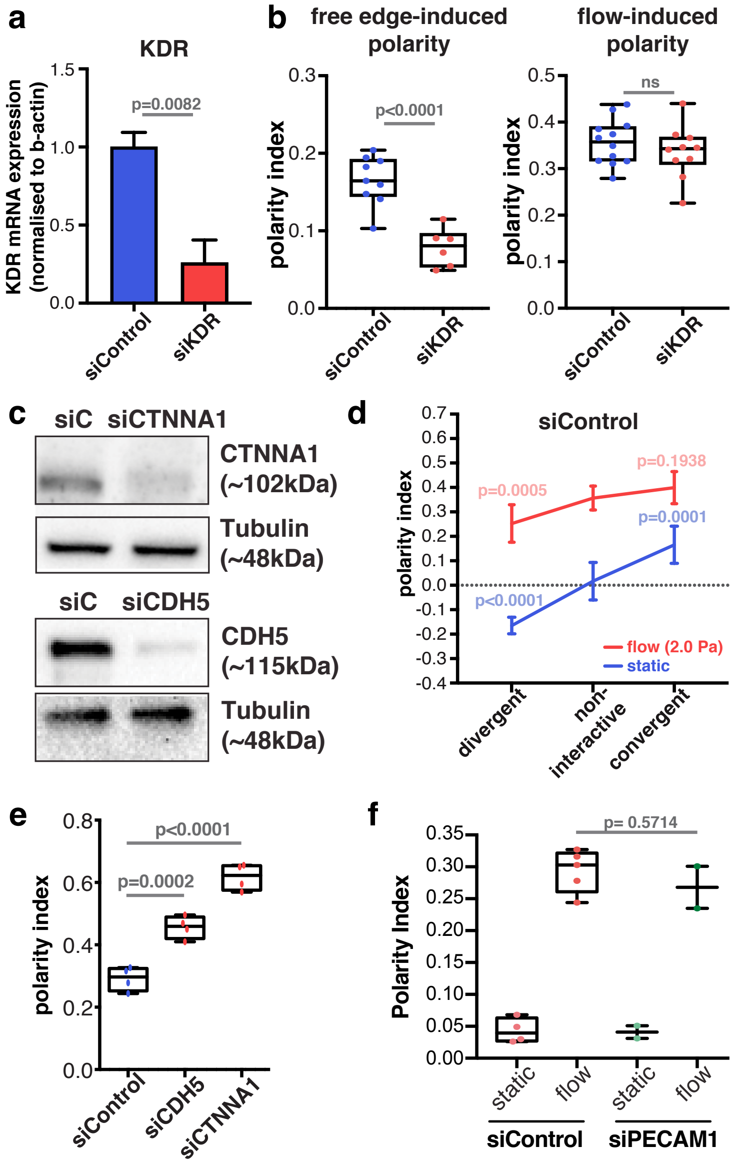


**Figure S9. Impairment of adherens junctions sensitizes cells to flow independently of the VE-cadherin-VEGFR2-PECAM1 complex.**
**a**, KDR mRNA expression levels in HUVECs treated with control siRNA (siControl) or siRNA targeting KDR (siKDR). N= 3 independent experiments. p-values from unpaired t test. **b,** Box plot of free edge-induce polarity index (left panel) and flow-induced polarity index (right panel) in siControl or siKDR HUVEC monolayers. N= 4 per condition; p-values from unpaired t test. **c**, Western blots for Ctnna1 and VE-cadherin upon depletion with siRNAs for siCTNNA1 and siCDH5, respectively. **d**, Polarity index for divergent, non-interactive and convergent regions in static (dark blue) or flow (dark red) conditions in HUVECs treated with siControl. N= 7 per condition; red p-values (flow 2.0 Pa) and blue p-values (static) correspond to multiple comparisons with non-interactive region one-way ANOVA corrected with Dunnett test. **e**, Box plot of flow-induced polarity index in siControl, siCTNNA1 or siCDH5 HUVEC monolayers. N= 4 per condition; p-values from unpaired t test. **f,** Boxplot for quantification in static or flow-stimulated HUVEC monolayers for siControl (N= 5) and siPECAM1 (N= 2). p-values from unpaired t test.

#### Figure S10

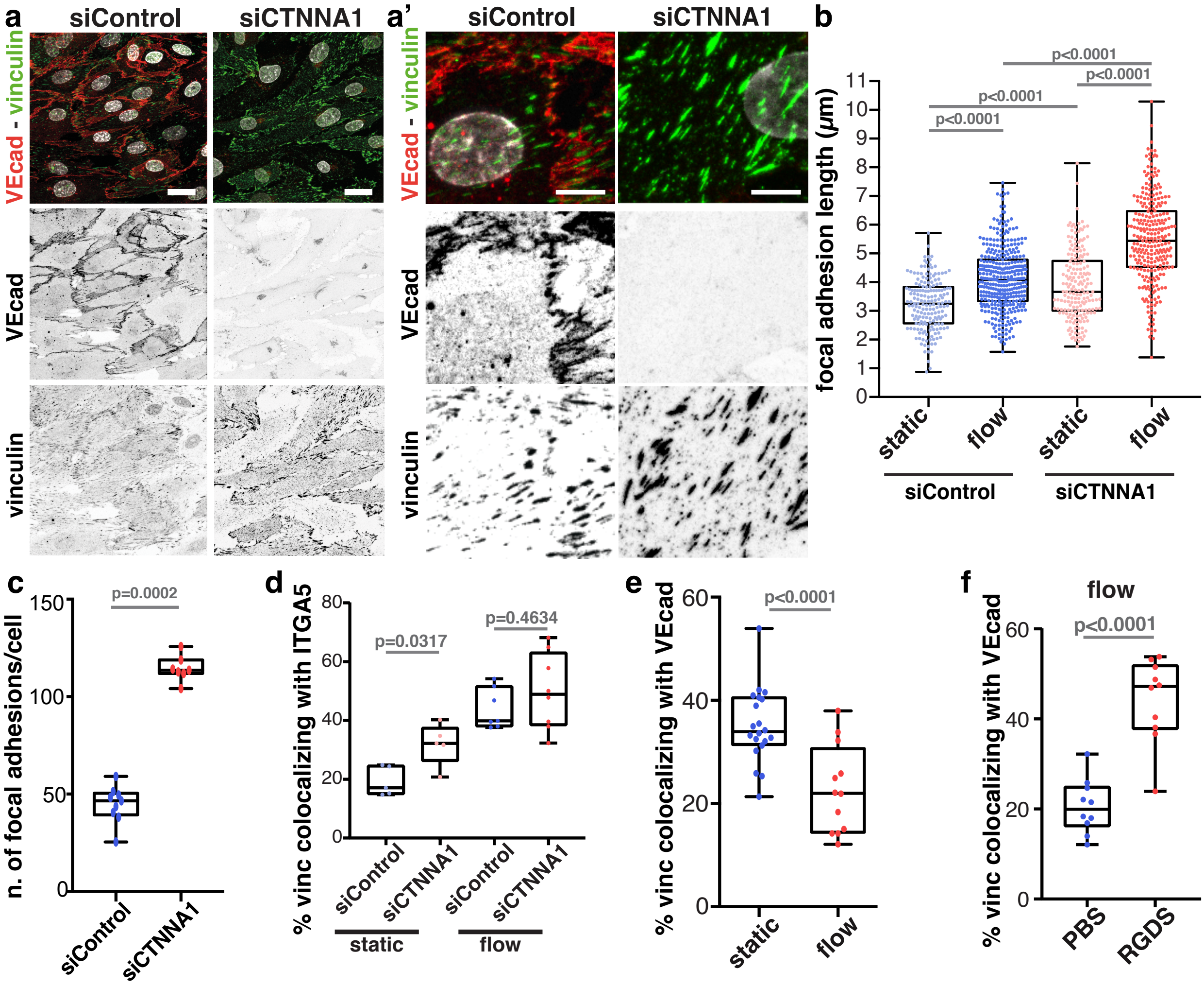


**Figure S10. Impairment of adherens junctions sensitizes cells to flow independently of the VE-cadherin-VEGFR2-PECAM1 complex.**
**a**, Representative images of flow-stimulated HUVEC monolayers for siControl and siCTNNA1, labelled for adherens junctions (VEcad, in red) and focal adhesions (vinculin, in green). Scale bar: 20um. A' zoomed image of A. **b**, Boxplot for quantification of focal adhesion length ($\mu$m) in static or flow-stimulated HUVEC monolayers treated with siControl or siCTNNA11. N> 300 per condition; p-values from unpaired t test. **c**, Box plot for quantification of number of focal adhesions in flow-stimulated siControl or siCtnna1 HUVEC monolayers. N= 8 per condition; p-values from Mann-Whitney test. **d**, Box plot for quantification of percentage of vinculin colocalizing with integrin alpha 5 (ITGA5) in static and flow-stimulated siControl or siCTNNA1 HUVEC monolayers. N= 5 per condition (static) and N= 7 per condition (flow); p-values from Mann-Whitney test. **e**, Box plot for quantification of percentage of vinculin colocalizing with VE-cadherin in flow-stimulated Control HUVEC monolayers. N= 20 (static) and N= 12 (flow); p-values from Mann-Whitney test. **f**, Box plot for quantification of percentage of vinculin colocalizing with VE-cadherin (VEcad) in flow-stimulated PBS- or RGDS-treated HUVEC monolayers. N= 10 per condition; p-values from Mann-Whitney test.

#### Figure S11

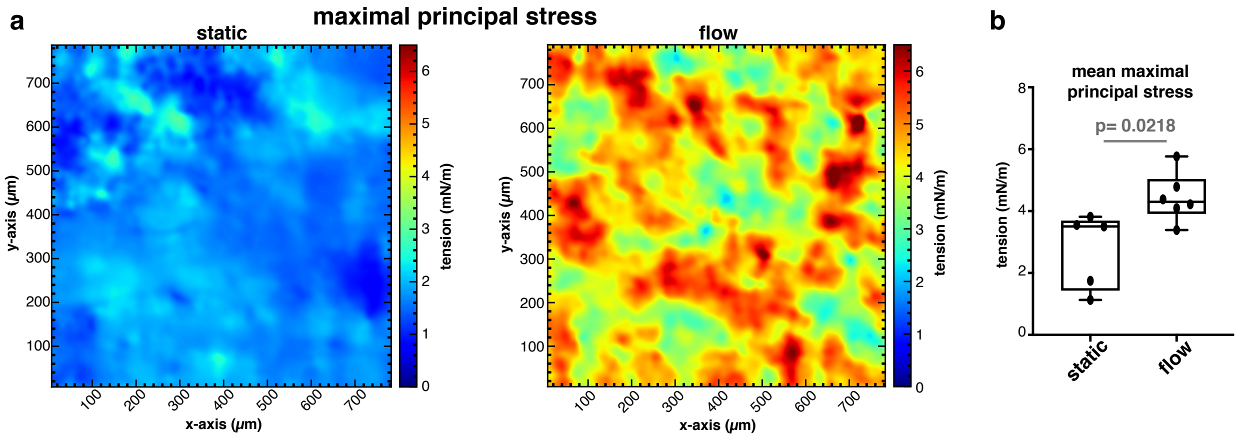


**Figure S11. Blood flow increases both traction forces and intercellular stresses.**
**a**, Representative images of maximal principal stress exerted by HUVECs in static or flow conditions obtained by monolayer stress microscopy. **b**, Average maximal principal stress exerted by HUVECs in static or flow conditions obtained by monolayer stress microscopy. N= 5 static and N= 6 flow; p-values from student t test.

#### Figure S12

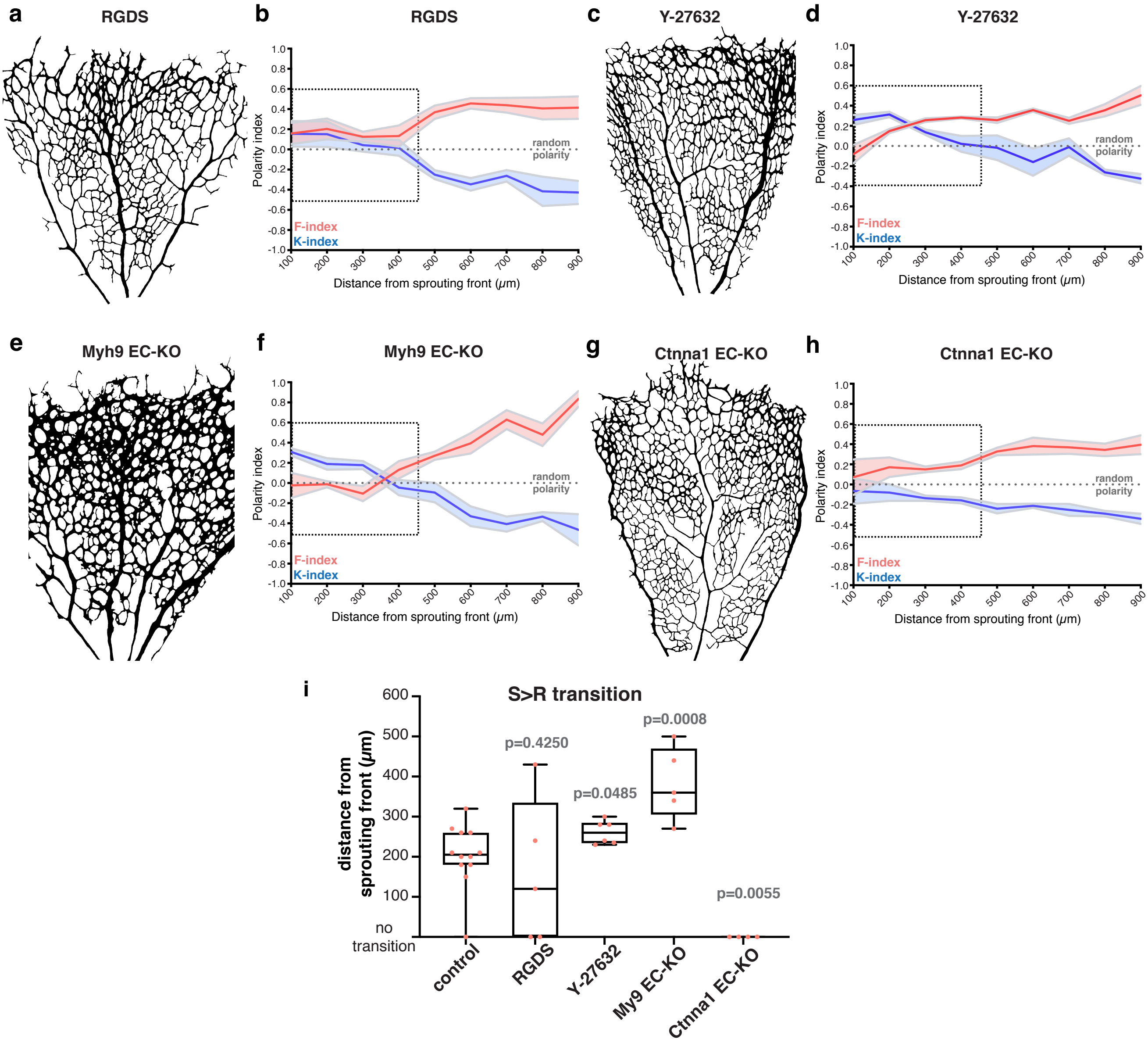


**Figure S12. Focal adhesion-mediated and adherens junction-mediated polarity regulates the S>R transition.**

**a**, Representative image of the vascular network following RGDS injection. **b**, Distribution of K-indexes (blue) and F-indexes (red) along the sprouting front to optic nerve axis of segmentation in RGDS injected mice. Solid line represents mean, and light area represents SEM. Dashed black line represents random polarity pattern. Dashed black rectangle highlights zoomed section showed in Fig.4. N= 5 arterial regions. **c**, Representative image of the vascular network following Y-27632 injection. **d**, Distribution of K-indexes (blue) and F-indexes (red) along the sprouting front to optic nerve axis of segmentation in Y-27632 injected mice. Solid line represents mean, and light area represents SEM. Dashed black line represents random polarity pattern. Dashed black rectangle highlights zoomed section showed in Fig.4. N= 5 arterial regions. **e**, Representative image of the vascular network of a Myh9 EC-KO mouse retina. **f**, Distribution of K-indexes (blue) and F-indexes (red) along the sprouting front to optic nerve axis of segmentation in Myh9 EC-KO mouse retinas. Solid line represents mean, and light area represents SEM. Dashed black line represents random polarity pattern. Dashed black rectangle highlights zoomed section showed in Fig.4. N= 5 arterial regions. **g**, Representative image of the vascular network of a Ctnna1 EC-KO mouse retina. **h**, Distribution of K-indexes (blue) and F-indexes (red) along the sprouting front to optic nerve axis of segmentation in Ctnna1 EC-KO mouse retinas. Solid line represents mean, and light area represents SEM. Dashed black line represents random polarity pattern. Dashed black rectangle highlights zoomed section showed in Fig.4. N= 4 arterial regions. **i**, Box plot for quantification of mean distance for the S>R transition in individual control, RGDS, Y-27632, Myh9 EC-KO and Ctnna1 EC-KO retinas. N= 11 for control; 5 for RGDS; 5 for Y-27632; 5 for Myh9 EC-KO; 4 for Ctnna1 EC-KO arterial regions. P-values from Mann-Whitney test between control and the corresponding group.
